# A host-pathogen tug-of-war over migrasome biogenesis in *Vibrio cholerae* infection

**DOI:** 10.64898/2026.11.16.766924

**Authors:** Peiyi Xia, Rui Wang, Ming Liu, Jiaxian Tang, Shu Zhang, Zhao Wang, Yuanpu Zhang, Shuo Yan, Hanhan Guo, Qi Xiang, Hao Wang, Hui Wang, Miao Tian, Yang Fu

## Abstract

Migrasomes are extracellular organelles implicated in cell communication and immune signaling, yet their induction by bacterial toxins and functional consequences during infection remains unknown. Here, we show that *Vibrio cholerae* infection induces non-canonical migracytosis in multiple cell types through the MARTX toxin. The actin cross-linking domain (ACD) promotes cell contraction and retraction fiber formation, while the α/β-hydrolase domain (ABH) collaborates with actin-polymerization machinery to support migrasome maturation. The Rho inactivation domain (RID) covalently modifies multiple Rho/Ras GTPases, and its absence yields oversized membrane blebs (large vesicles) enriched in endoplasmic reticulum, lipid droplets, and pro-inflammatory chemokines compared with wild-type migrasomes. In an infant mouse model, RID-deficient *V. cholerae* exhibits impaired intestinal colonization and elevated host chemokine transcription. These findings establish that MARTX-driven migrasome formation is not merely a byproduct of cytoskeleton damage, but represents a coordinated, domain-specific reprogramming of host membrane dynamics with implications for host-pathogen interplay.

## Introduction

Migrasomes are recently discovered extracellular organelles that form on retraction fibers during cell migration (Ma et al., 2015). The biogenesis of migrasomes, a process termed migracytosis, is strictly orchestrated in three stages (Jiang et al., 2025): nucleation, initiated by sphingomyelin synthase 2 (SMS2) (Liang et al., 2023); maturation, involving the recruitment of ITGA5 via PI(4,5)P2 and Rab35 (Wu et al., 2017; Zhen and Stenmark, 2023); and stabilization, characterized by the assembly of tetraspanin-cholesterol macrodomains (Dharan et al., 2022; Dharan et al., 2023; Huang et al., 2019). During biogenesis, specific cellular cargo is selectively sorted into migrasomes via Rab protein-mediated trafficking (Jiao et al., 2024a). These sequestered contents enable migrasomes to participate in a diverse array of biological processes. For example, migrasomes provide regional cues for organ morphogenesis via CXCL12 signaling (Jiang et al., 2019), the maintenance of cellular homeostasis through “mitocytosis”—the disposal of damaged mitochondria (Jiao et al., 2021) and maintain haemostasis system by migrasomes produced from neutrophils (Jiang et al., 2024).

Beyond these physiological roles, migrasomes are increasingly recognized as critical players in infection and immunity. Recent reports indicate that viruses can induce migracytosis and exploit migrasomes for dissemination (Huang et al., 2025; Zhang et al., 2022), whereas the inhibition of migrasome biogenesis effectively blocks viral spread (Hou et al., 2026; Xu et al., 2025). Monocytes stimulated by lipopolysaccharide (LPS) have been shown to load cytokines into migrasomes for systemic signal transmission through the circulatory system (Jiao et al., 2024b). Recent evidence shows that *Clostridioides difficile* TcdB, a Large Clostridial Toxins (LCTs) family protein that inactivate RHO/RAS GTPases through glycosylation (Orrell and Melnyk, 2021), have been shown to trigger “non-canonical migracytosis” to exacerbate early immune responses (Li et al., 2024).

Bacterial toxins that target the Rho GTPases represent a major class of virulence factors, with diverse families disrupting actin dynamics or cell-cell junctions to create permissive niches for colonization (Aktories, 2011; Voth and Ballard, 2005). For example, TcdB disrupts the actin cytoskeleton resulting in cell rounding (Orrell and Melnyk, 2021), the retraction fiber foundationally formed based on the non-migrate cell deformation. Beside the LCTs, bacterial pathogens secrect toxin diversely remodel host actin cytoskeleton (Popoff, 2024). The *Vibrio* genus deploys such toxins, the Multifunctional Autoprocessing Repeats-in-Toxin (MARTX), which combines multiple enzymatic domains to simultaneously manipulate actin polymerization, hijack GTPases signaling and so on (Olivier et al., 2009). Yet whether these distinct strategies similarly converge on migrasome biogenesis remains unknown.

In *Vibrio cholerae*, the MARTX toxin is a large, modular protein that delivers multiple enzymatic domains into the host cytoplasm (Satchell, 2015). Autoprocessing is mediated by an internal Cysteine Protease Domain (CPD) activated by host-cell inositol hexakisphosphate (InsP6), which triggers the release of effector cargo (Prochazkova et al., 2009). The best-characterized domains include the Actin Cross-linking Domain (ACD), which covalently crosslinks G-actin to disrupt the cytoskeleton (Kudryashov et al., 2008a; Kudryashov et al., 2008b); the α/β-Hydrolase (ABH) domain, a phospholipase targeting PI3P to modulate autophagy and membrane trafficking (Agarwal et al., 2015); and the Rho Inactivation Domain (RID), which catalyzes Nε-fatty acylation of RHO and RAS GTPases (Zhou et al., 2017), to hijack cellular signaling (Geissler et al., 2010). In *Vibrio vulnificus*, the MARTX toxin consistently retains the RID and ABH domains in highly virulent clinical isolates (Kwak et al., 2011). *In vivo* infection models have demonstrated that these domains play a critical role in modulating host survival and the inflammatory response (Woida and Satchell, 2020). Notably, while individual MARTX domains have been studied in isolation, how ABH, ACD and RID act in concert to remodel the host cytoskeleton and whether this concerted action shapes migrasome biogenesis remains unknown.

Here, we investigated whether and how *V. cholerae* infection triggers migrasome biogenesis and the functional consequences of toxin-induced migrasome formation during infection. Using a combination of live-cell imaging, genetic manipulation of MARTX domains, click chemistry, quantitative proteomics, cryo-electron tomography, and an infant mouse infection model, we dissected the contributions of individual MARTX effector domains to migrasome formation and cargo composition. Our findings establish MARTX-driven migrasome biogenesis as a coordinated, domain-specific reprogramming of host membrane dynamics and highlight the formation and regulation of migrasomes as a potential nexus in host-pathogen interplay.

## Result

### *Vibrio* Infection Induces Migrasome Formation in Multiple Cell Types

To investigate whether bacterial infection triggers migrasome biogenesis, we first examined the morphological changes of Caco-2 cells upon challenge with various *Vibrio* species. Confocal microscopy of WGA-stained cells revealed that within 4 hours of infection, cells exhibited dramatic morphological alterations characterized by the formation of retraction fibers (RFs) emanating from the cell body, with numerous vesicular structures budding along these fibers (Figure S1A). Quantification showed that these extracellular vesicles ranged from 0.5 to 3 μm in diameter (Figure S1B), consistent with the reported size distribution of migrasomes (Ma et al., 2015).

To verify the identity of these vesicles, we established stable Caco-2 cell lines expressing established migrasome markers, including TSPAN4-mCherry and ITGA5-eGFP. Live cell confocal imaging following infection with GFP-labeled *Vibrio cholerae* C6706 revealed that the markers prominently localized to both RFs and extracellular vesicles (Figure 1A). To functionally validate migrasome formation, we employed two complementary approaches. First, CRISPR-Cas9 mediated knockout of *ITGA5*, an essential protein for migrasome biogenesis, abolished the formation of migrasomes upon bacterial infection (Figure 1B, 1D and Figure S1E). The dual-sgRNAs knockout efficiency was confirmed by genomic PCR and western blot assay (Figure S1C-D). Second, pharmacological inhibition of sphingomyelin, a critical lipid component for migrasome formation, similarly prevented migrasome generation (Figure 1C-D). Together, these genetic and pharmacological manipulations establish that *V. cholerae* infection induces bona fide migrasome formation.

**Figure 1.**
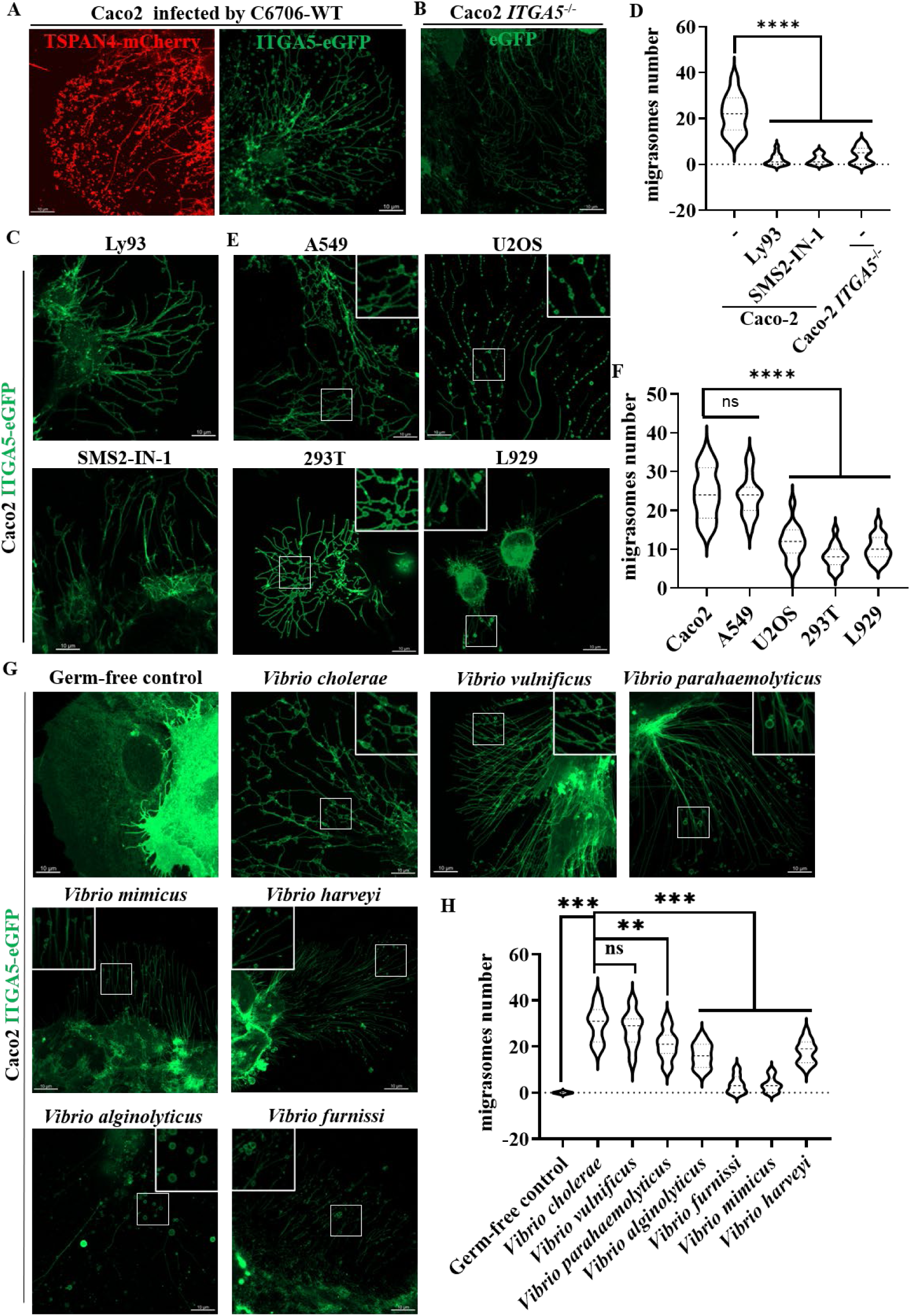
Vibrio infection induces passive migration and migrasome biogenesis. (A) Representative images of Caco2 cells stably expressing TSPAN4-mCherry and ITGA5-eGFP following infection with wild type *V. cholerae* C6706 (C6706-WT). (B) Confocal images Caco2 ITGA5^−/−^stably express eGFP was infected by C6706-WT. (C) Confocal images *V. cholerae* infects Caco-2 cells express ITGA5-eGFP. The Caco-2 cells were treated with Ly93 or SMS2-IN-1 1 hour in advance. (D) Quantification of the numbers of migrasome from (A-C) are measured as mean ± SEM; n=15. P values were calculated using one-way ANOVA followed by Dunnett’s test. (E) A549, U2OS, 293T and L929 cell lines stably express ITGA5-eGFP were infected by C6706-WT. (F) Quantification of the numbers of migrasome from (E) are measured as mean ± SEM; n=15. P values were calculated using one-way ANOVA followed by Dunnett’s test. (G) *Vibrio*-induced migrasome formation in ITGA5-eGFP Caco2 cells across different Vibrio species. (H) Quantification of the numbers of migrasome from (G) are measured as mean ± SEM; n=15. P values were calculated using one-way ANOVA followed by Dunnett’s test.

We next asked whether this phenomenon is restricted to Caco-2 cells or specific to *V. cholerae*. Strikingly, ITGA5-eGFP marked migrasomes were observed across diverse cell lines upon *V. cholerae* infection (Figure 1E-F). Furthermore, multiple pathogenic *Vibrio* species also triggered robust migrasome formation (Figure 1G). Quantitative analysis confirmed migrasome production across all tested conditions, albeit with some variation in efficiency (Figure 1H). Moreover, Caco-2 infect with *E. coli* MG1655 would not undergo migracytosis (Figure S1F). These results demonstrate that *Vibrio*-induced migrasome formation is not isolated phenomenon but rather a widespread cellular response to infection.

### MARTX Toxin is the Essential Factor Driving non-canonical migracytosis

To distinguish whether migrasome formation requires direct bacterial contact, we performed time-lapse imaging of TSPAN4-mCherry expressing Caco-2 cells challenged with GFP-labeled *V. cholerae*. Strikingly, cell deformed, retracted and generated migrasomes before visible bacterial contact (Figure S2A), indicating that a diffusible secreted factor drives this process. To identify this factor, we screened isogenic mutants lacking secretion toxins: Δ*vipA*, Δ*hlyA/hapA*, Δ*ctxAB* or Δ*rtxCPD* (rendering MARTX non-functional). Only cells infected by C6706-Δ*rtxCPD* failed to induce migrasomes, while all other mutants produced them at wild-type levels (Figure 2A-B). These results establish the MARTX toxin as the unique and essential bacterial determinant of migrasome biogenesis.

**Figure 2.**
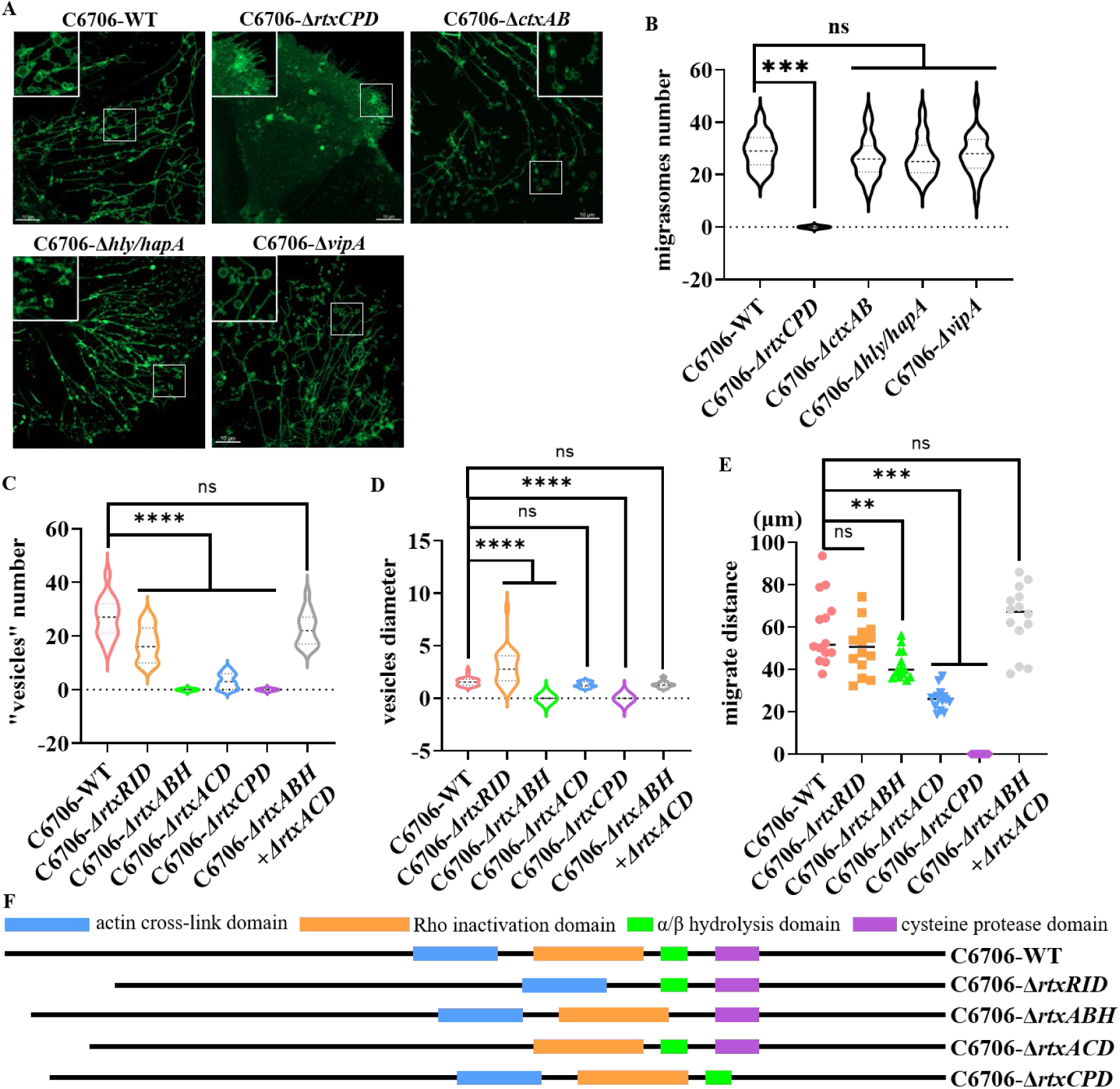
The MARTX toxin is the primary driver of *Vibrio cholerae* induced non-canonical migracytosis. (A) Representative confocal images showing the induction of migrasome formation in Caco2 cells following infection with various *V. cholerae* secretion mutants. (B) Quantification of the numbers of migrasome from (A) are measured as mean ± SEM; n=25. P values were calculated using one-way ANOVA followed by Dunnett’s test. (C) – (E) Quantification of the numbers of “vesicles” (C), the “vesicles” dimeter (D) and the cell migrate distance (E) from (fig. S2B-C) are measured as mean ± SEM; n=20. P values were calculated using one-way ANOVA followed by Dunnett’s test. (F) Schematic model of defective *V. cholerae* C6706.

MARTX toxin comprises multiple enzymic domains that auto-process via an internal cysteine protease. To dissect their individual contributions to migrasome formation, we infected cells with mutants lacking ABH (Δ*rtxABH*), ACD (Δ*rtxACD*) or RID (Δ*rtxRID*), alongside the inactive Δ*rtxCPD* control (Figure 2F). Caco-2 cells infected with the C6706-Δ*rtxCPD* abolished all migrasome formation, infected by C6706-Δ*rtxABH* produced minimal migrasomes despite residual cell migration, infected by C6706-Δ*rtxACD* showed reduced cell migration distance but residual migrasome-like vesicles, infected by C6706-Δ*rtxRID* migrated normally but more easily generated larger, irregular membrane sheets rather than typical migrasomes (Figure 2C-E, Figure S2B). Notably, when C6706-Δ*rtxABH* and C6706-Δ*rtxACD* co-infected with Caco-2 cells, the cells phenomenon would be rescued as infected with wild-type C6706 (Figure 2C-E, and Figure S2B(i)-C).

### MARTX-driven migrasome benefits for *V. cholerae* adhesion

*V. cholerae* depends on MARTX toxin to disrupt epithelial barriers during intestinal colonization. To determine whether the observed domain-specific effects on host cells translate into altered bacterial fitness, we performed a qPCR assay and adhesion assay. After being infected by the mutant genotype bacteria for 2 or 3 hours, we collected the infected cells after washing twice with PBS. At 2 hours post-infection, we found that the CFU did not exhibit a difference whereas the *tcpA* transcription level in C6706-WT showed a range from 3 to 9 times higher than the other mutant C6706. At 3 hours post infection, the fold change in transcription level was reduced while the CFU reflected differences: C6706-Δ*rtxCPD* exhibited a ∼70% reduction in bacterial attachment versus WT, while C6706-Δ*rtxRID* decreased only about 25%, C6706-Δ*rtxABH* and C6706-Δ*rtxACD* showed ∼50-60% reductions (Figure 3A-B). Meanwhile, we examined whether migrasome could influence the bacteria adhesion ability. At 2 hours post infection, we found that the C6706-WT *tcpA* transcription level in infecting Caco-2 WT was 2 times higher than in infecting Caco-2 *ITGA5*^−/−^. As a consequence, the adhesion ability of C6706-WT showed ∼50% reduction in migrasome biogenesis defective Caco-2 cells at 3 hours post infection (Figure 3C-D).

**Figure 3.**
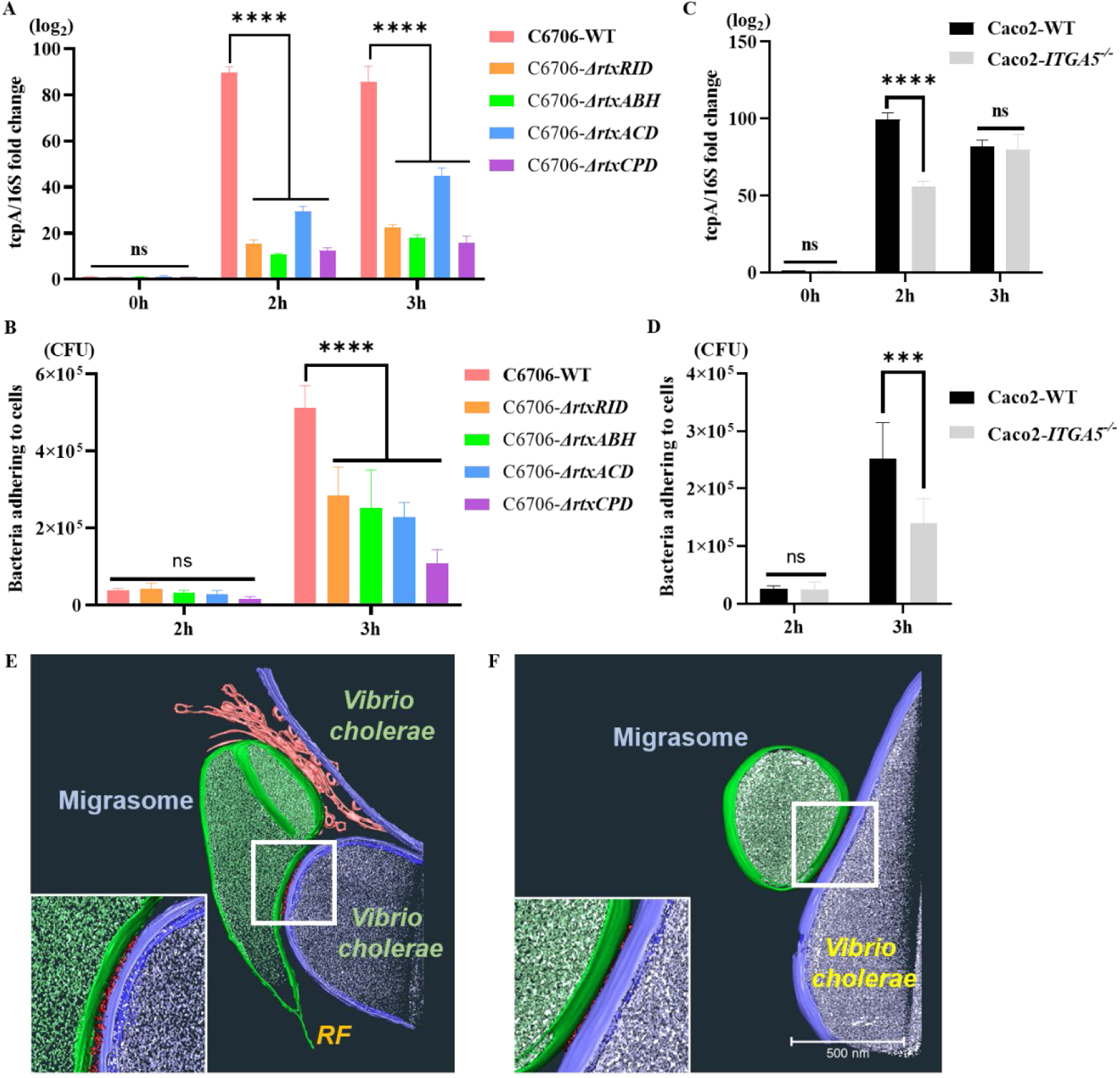
*Vibrio cholerae* directly interact with migrasomes. (A) and (B) qPCR reflect *tcpA*/16S transcript relative fold change of C6706 at two or three hours post infection. (A) the transcript relative fold change in various genotype C6706. (C) the transcript relative fold change of C6706-WT during infecting Caco-2 WT or Caco-2 *ITGA5*^−/−^cells. (B) and (D) CFU reflect the number of *V. cholerae* adhesion to Caco-2 cells. (B) various genotype C6706 adhesion to Caco-2 cells. (D) C6706-WT adhesion to Caco-2 WT or Caco-2 *ITGA5*^−/−^cells. (E) and (F) tomography reflects the interaction between migrasomes (green) and *V. cholerae* (blue) via potential proteins indicated by densities (red). (E) *in vivo* condition, migrasomes are generated from live cells. (F) *in vitro* condition, migrasomes are purified from infected cells.

Meanwhile, we used cryo-electron tomography (Cryo-ET) to investigate whether *V. cholerae*directly interacts with migrasomes. Caco-2 cells were grown on grids and infected with C6706-WT once they reached ∼50% confluence. After 3 to 4 hours of infection, the cells were plunge-frozen and subjected to cryo-ET using a 300 kV cryo-electron microscope. We observed electron-dense material (red) between C6706-WT (blue) and migrasomes (green), which may represent proteins or glycans that mediate bacteria-migrasome interactions (Figure 3E). To exclude the possibility that the observed interactions were merely due to random collisions, we purified migraomes from infected Caco-2 cells and incubated them with C6706-WT for 2 hours before Cryo-ET sample preparation. Tomographic reconstruction showed that C6706-WT still interacted with purified migrasomes under these cell-free conditions, indicating that the interaction between *V. cholerae* and migrasomes is being mediated by specific surface components (Figure 3F).

### ABH and ACD Collaborate to Drive Actin Polymerization for Migracytosis

Given that C6706-Δ*rtxABH* and C6706-Δ*rtxACD* both impaired migrasome formation and bacterial adhesion, we examined the actin cytoskeleton as a potential connection. SDS-PAGE analysis of actin oligomerization states revealed that C6706-WT infection triggered progressive actin polymerization. Unexpectedly, we found C6706-Δ*rtxABH* infection barely caused actin oligomerization (Figure S3A). Due to the difficulty to complement the ABH domain in C6706-Δ*rtxABH*, we came up with a method to simply mimic the complementary experiment: infect Caco-2 cells with the mixed C6706-Δ*rtxABH* and C6706-Δ*rtxACD*. As mentioned above, in this condition, we observed some cells undergo migracytosis, the same as being infected by C6706-WT, which cannot happen in either C6706-Δ*rtxABH* or C6706-Δ*rtxACD* infection (Figure 2C-E, Figure S2B-C).

Thus, we focus on trying to clarify how ABH could influence ACD function in Caco-2 cells. Taking actin polymerization as a bridge between MARTX toxin domains and migracytosis, we used time-lapse confocal microscopy in Caco-2 cells co-expressing ITGA5-eGFP and lifeact-miRFP to investigate the actin movement in migracytosis. Combined with a time-course assay, we collect cells every hour post-infection by C6706-WT or C6706-Δ*rtxABH* and utilize immunoblot to investigate the degree of actin polymerization. We observed that during C6706-WT infection, cells began retracting within the first hour with actin dimerized; in the subsequent hour, actin oligomerized into higher-order polymers that migrated outward along retraction fibers, ultimately seeding migrasome growth at discrete positions (Figure 4A-B, Figure S3C). In contrast, C6706-Δ*rtxABH* infected cells also underwent retraction and produced retraction fibers, but the process was markedly prolonged and even after three hours no migrasomes had formed, with actin remaining predominantly as dimers and monomers (Figure 4C-D, Figure S4D). Further, we disrupted F-actin alone as a control. After treating uninfected Caco-2 cells with cytochalasin D, we observed cells contraction and retraction fiber formation due to F-actin disassembly. Yet despite the presence of extensive retraction fibers, no migrasomes were observed (Figure 4F). This dissociation between cell retraction and migrasome formation indicated that during *V. cholerae* infection, actin polymerization can be a way for migracytosis.

**Figure 4.**
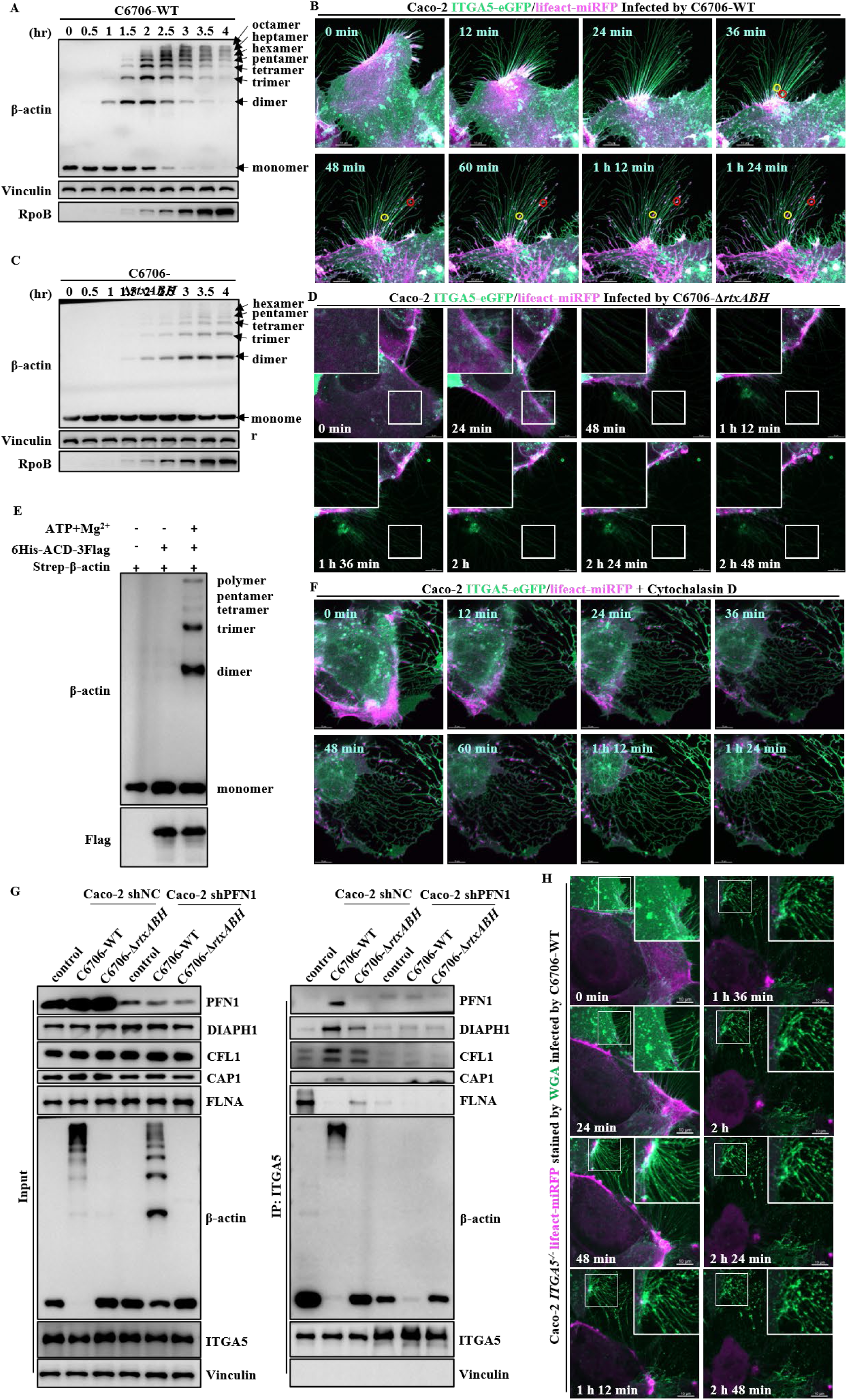
Hierarchical actin polymerization drives cell contraction and migrasome biogenesis. (A and C) Time-course Western blot analysis of actin polymerization in Caco-2 cells during infection with (A) C6706-WT or (C) the C6706-Δ*rtxABH*. Samples were collected at 30-minute intervals post-infection (MOI = 10). (B and D) Representative time-lapse confocal images of Caco-2 cells stably expressing ITGA5-eGFP and Lifeact-miRFP (actin marker) following infection with (B) C6706-WT or (D) C6707-Δ*rtxABH*. (E) *In vitro* biochemical assay demonstrating that the MARTX ACD domain is sufficient to induce low-order β-actin crosslinking in the presence of Mg^2+^ and ATP. (F) time lapse confocal imaging of Caco-2 ITGA5-eGFP/Lifeact-miRFP treated with 5 μM Cytochalasin D. (G) Co-immunoprecipitation assay of Caco-2 shNC or Caco-2 shPFN1 infected by C6706-WT or C6706-Δ*rtxABH*, germ-free panel as negative control. (H) Time-lapse confocal images WGA stained Caco-2 *ITGA5^−/−^*cells stably expressing Lifeact-miRFP infected with the C6706-WT.

To identify these cofactors, we performed immunoprecipitation from C6706-WT infected cells followed by mass spectrometry. This analysis revealed a cohort of actin-polymerization machinery components co-enriched with actin during infection, including PFN1, DIAPH1, PFN2, VASP, CAP1, CFL1 and FLNA (Figure S4A-B). These proteins are established regulators of G-actin incorporation into filaments and of F-actin linkage to integrin adhesion complexes. To determine whether these cofactors participate in MARTX toxin-mediated actin cross-linking, we individually knocked down each candidate by shRNA (Figure S4C). Under C6706-WT infection, PFN1-knockdown Caco-2 cells showed markedly impaired actin polymerization compared to shNC control cells within the same time (Figure 4G, left). We next performed co-immunoprecipitation experiments to investigate whether PFN1 knockdown disrupts the interaction between actin and ITGA5. Using ITGA5 as bait, we found that in shNC Caco-2 cells, ITGA5 co-precipitated monomeric or polymerized actin across all conditions (uninfected controls, C6706-WT and C6706-Δ*rtxABH* infection). However, PFN1, DIAPH1, CAP1 and CFL1 were co-precipitated with ITGA5 only during C6706-WT infection (Figure 4G). Upon PFN1 knockdown, ITGA5 no longer co-precipitated polymerized actin from C6706-WT infected cells, yet monomeric actin from uninfected or C6706-Δ*rtxABH* infected cells remained detectable (Figure 4G). Consistently, PFN1 knockdown also abolished the co-precipitation of DIAPH1, CAP1 and CFL1 with ITGA5 (Figure 4G). To examine whether PFN1 contributes to migrasome formation, we generated a stable Caco-2 cell line co-expressing TSPAN4-mCherry and lifeact-miRFP with PFN1 knockdown. Long-term live imaging revealed that these cells required a longer time to undergo morphological deformation, and notably, failed to produce abundant migrasomes even at 2.5 hours post-infection (Figure S4D-F). In ITGA5-knockout cells, polymerized actin accumulated within the cell body but failed to transport outward along retraction fibers, resulting in near-complete loss of migrasome formation (Figure 4H).

### RID Restricts Membrane Blebbing and Shapes Migrasome Morphology Through Covalent Modification of RHO/RAS GTPases

While the ABH and ACD domains collaboratively generate the actin polymers necessary for migrasome biogenesis, the RID domain plays a distinct yet equally critical role in determining the ultimate morphology and cargo composition of the extracellular vesicles. Live-cell confocal imaging of Caco-2 cells stably expressing ITGA5-eGFP and lifeact-miRFP infected with the C6706-Δ*rtxRID* revealed a striking phenotypic divergence from WT infection. Rather than observing the outward transport of polymerized actin along retraction fibers that culminates in migrasome formation, cells infected with the RID deficient strain displayed extensive membrane blebbing at sites of actin polymerization (Figure S5A). These membrane sheets appeared to be physically trapped by intact F-actin networks as the cell body retracted, suggesting that the absence of RID permits F-actin to retain its membrane-anchoring capacity during cell contraction. To test whether intact F-actin was indeed sufficient to trap membranes, we treated cells with cytochalasin D to induce F-actin depolymerization and observed that membrane blebbing was significantly attenuated, confirming that robust F-actin assembly is required for the large membrane sheet phenotype (Figure S5B). Notably, we detected the ITGA5 content in migrasomes and the blebbed membranes, we found that ITGA5 seems not necessary in the large membrane sheets formation (Figure 5E). Further, ITGA5-knockout Caco-2 cells infected with the C6706-Δ*rtxRID* still produced large membrane sheets that eventually resolved into spherical vesicles at later time points, demonstrating that this process occurs independently of ITGA5-mediated actin outward transport (Figure S5C). Quantitative morphological analysis by confocal microscopy and negative-stain electron microscopy revealed dramatic differences between vesicles produced under C6706-WT versus C6706-Δ*rtxRID* infection conditions (Figure 5A-B). C6706-WT infection generated uniformly spherical migrasomes with an average diameter from 0.5 μm to 3 μm, whereas C6706-Δ*rtxRID* infection yielded irregularly shaped membrane sheets then forming spherical vesicles of varying sizes (Figure 2D).

**Figure 5.**
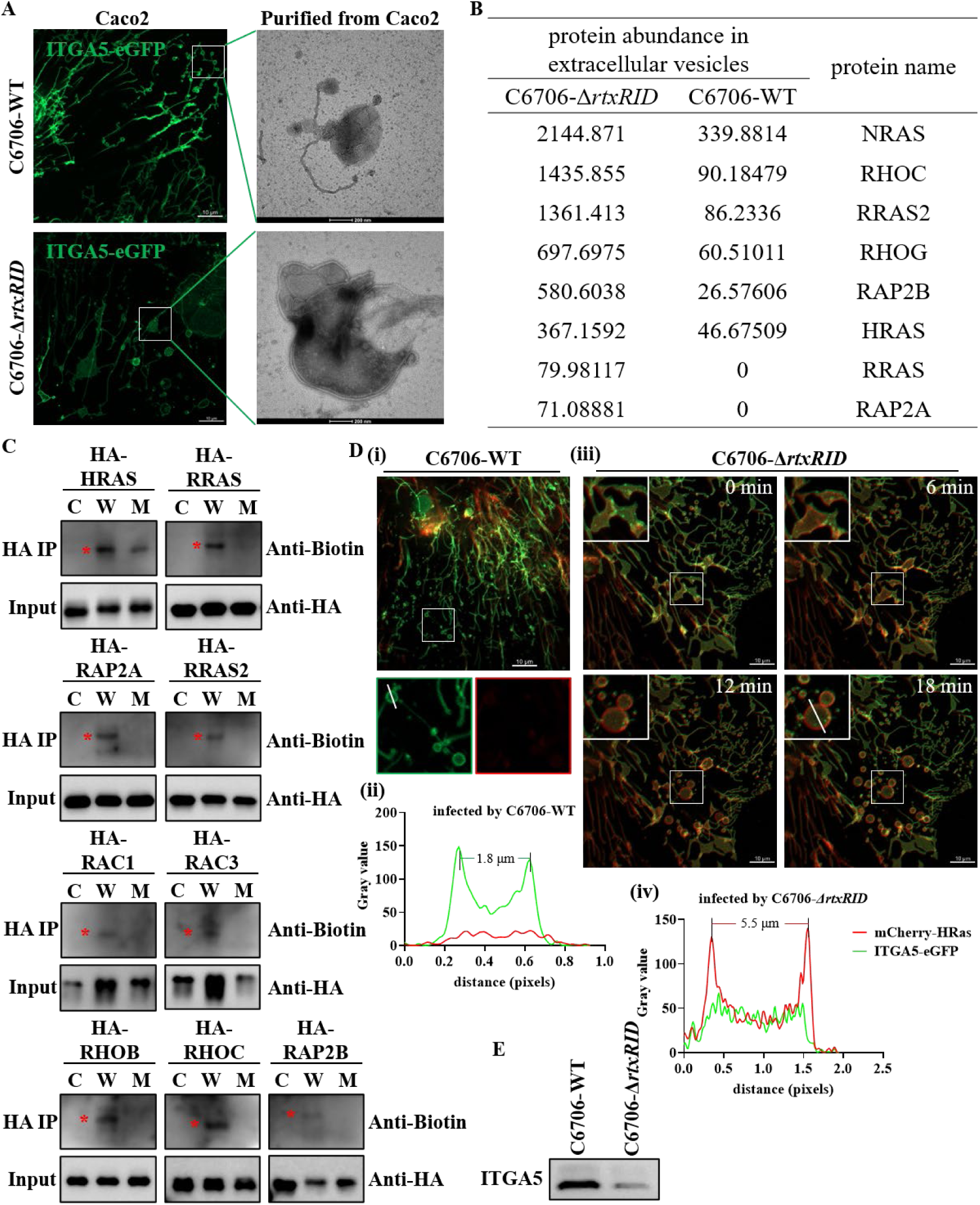
H-Ras recruitment governs the morphological divergence of *V. cholerae*-induced vesicles. A) Comparative morphological analysis of C6706-WT migrasomes and the enlarged vesicles induced by C6706-Δ*rtxRID*. Representative live-cell confocal micrographs (left) and transmission electron microscopy (TEM) images (right) of purified vesicles are shown. (B) Quantitative mass spectrometry (MS) analysis revealing the differential enrichment of RHO/RAS GTPases in C6706-Δ*rtxRID* induced vesicles compared to C6706-WT migrasomes. (C) click chemistry validate the RHO/RAS GTPases modified by RID. (D) Localization and recruitment dynamics of H-Ras. Caco2 cells were triple-transduced to stably express ITGA5-eGFP, Lifeact-miRFP, and mCherry-H-Ras. Cells were challenged with either (Di) C6706-WT or (Diii) the C6706-Δ*rtxRID*. (Dii, and Div) Fluorescence intensity profiles (line-scans) generated using ImageJ along the indicated white line. (E) Western blot analysis of purified migrasomes and C6706-*ΔrtxRID* induced large vesicles, characterizing the protein content of the ITGA5.

To investigate the molecular mechanism underlying RID-dependent vesicle morphogenesis, we employed click chemistry to systematically profile which Rho/Ras family GTPases are covalently modified by RID during infection. Using HA-tagged GTPases and biotin-based detection in a RID-dependent manner: H-Ras, R-Ras, R-Ras2, Rap2A, Rap2B, Rac1, Rac3, RhoB and RhoC (Figure 5C). Notably, this modification was abolished in C6706-Δ*rtxRID* infection, confirming the specificity of RID enzymatic activity. By labeling the GTPases that can be modified by RID with mCherry, we trying to track the function of the GTPases in membrane morphology control during cell retraction and migration. We applied time-lapse live cell imaging assay by infect the Caco-2 cell which stably expressed ITGA5-eGFP and mCherry labeled GTPases. Among the RID-modified GTPases, we found only H-Ras uncovered a complementary role in vesicle morphogenesis. Under C6706-WT infection, HRAS signal is barely detectable in migrasome whereas ITGA5 signals concentrated specifically at migrasome budding sites, with intensity profiles showing sharp peaks corresponding to individual migrasomes (∼1.8 μm width) (Figure 5D (i)-(ii)). Conversely, H-Ras signal distributes on the membrane under C6706-Δ*rtxRID* infection, the signal intensity increased steadily until resolve the membrane sheets into spherical vesicles. The intensity profiles of H-Ras showing sharp peaks corresponding to the spherical vesicles about 5.5 μm (Figure 5D (iii)-(iv)).

Given that several Rho/Ras family GTPases are modified by RID in Caco-2 cells, we sought to understand why membrane rupture occurs during cell retraction in cells infected with C6706-Δ*rtxRID*. As with H-Ras, we generated mCherry-tagged versions of candidate GTPases and tracked their subcellular localization by time-lapse confocal microscopy. We observed that Rac3 may play a role in large vesicle stabilization (Figure S6A), whereas RhoG and RhoC may promote membrane rupture (Figure S6B-C), even though RhoG is not a substrate of RID (Figure 5C).

### Migrasomes and Large Vesicles Function as a Platform for Extracellular Cargo Release and Bacterial interaction

The morphological differences between C6706-WT derived migrasomes and C6706-Δ*rtxRID* induced membrane blebs raised a critical question: do these structurally distinct vesicles differ in their subcellular composition and functional cargo? To systematically compare the content profiles, we purified extracellular vesicles from Caco-2 cells infected with either C6706-WT or C6706-Δ*rtxRID* and performed quantitative mass spectrometry. Subcellular localization analysis revealed markedly different distributions of vesicle constituents. C6706-WT derived migrasomes were predominantly enriched in cytoplasmic proteins, extracellular matrix and secretory vesicle components, with notably minimal representation of endoplasmic reticulum (ER) and lipid droplet (LD) markers. In contrast, C6706-Δ*rtxRID* derived vesicles exhibited a significantly altered composition, with reduced extracellular matrix components and a relative enrichment of ER-associated proteins, suggesting that the absence of RID permits bulk membrane capture that entraps diverse organellar contents during aberrant vesiculation (Figure 6A-B).

**Figure 6.**
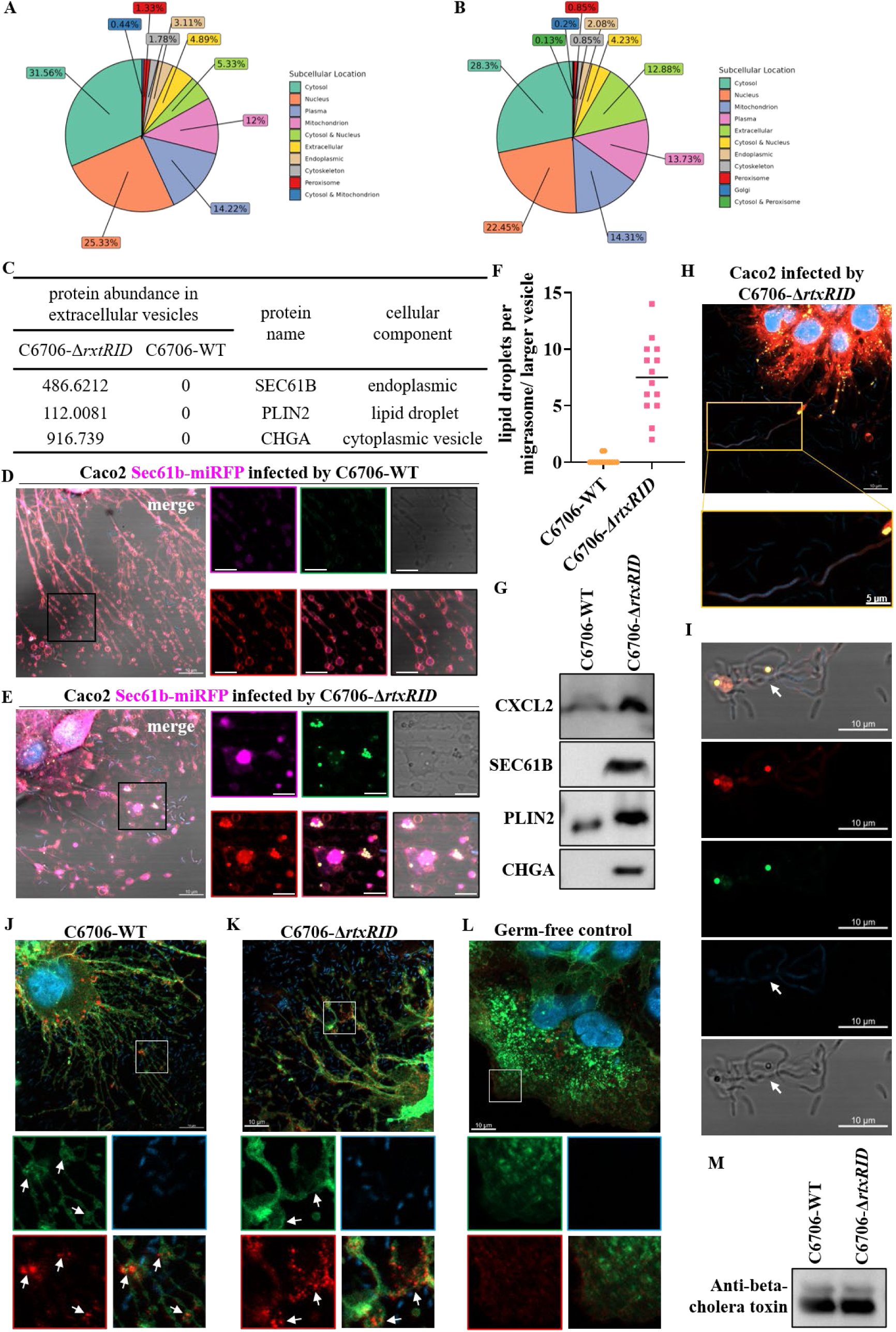
MARTX-mediated regulation of migrasome biogenesis determines subcellular cargo and chemokine sequestration. (A) and (B) subcellular location analysis of proteomics data. (A) analysis of migrasomes. (B) analysis of large vesicles. (C) Quantitative mass spectrometry (MS) analysis revealing the enrichment of endoplasmic reticulum (ER) and lipid droplet (LD) proteins in C6706-*ΔrtxRID* induced vesicles, or in C6706-WT induced migrasomes. (D) and (E) Representative live-cell confocal images Caco-2 cells stably express Sec61b-miRFP (ER marker) and stained by Nile Red infected either with (D) C6706-WT or (E) C6706-*ΔrtxRID*. Under differential excitation (556 nm for neutral lipids/membranes and 488 nm for lipid droplets), plasma membranes are visualized in red and lipid droplets in yellow. (F) lipid droplets number in each migrasome or larger vesicle was statistical quantified manually (n=14). (G) Western blot analysis of purified migrasomes and C6706-*ΔrtxRID* induced large vesicles, characterizing the protein content of the chemokine CXCL2, endoplasmic reticulum marker SEC61B, the lipid droplet marker PLIN2 and an antimicrobial protein CHGA. (H)-(I) Confocal images revealing elongated C6706-*ΔrtxRID* interact with LDs. (H) LDs in large vesicles were marked by Nile Red in yellow and C6706-*ΔrtxRID* were stained by Hoechst. (I) isolated LDs were marked by Nile Red in yellow and C6706-*ΔrtxRID* were stained by Hoechst. (J) - (L) immunofluorescence validation of CXCL2 localization and distribution. Representative images are shown for (J) C6706-WT infection, (K) C6706-*ΔrtxRID* infection and (L) an uninfected (germ-free) control. (M) Western blot analysis of purified migrasomes and C6706-*ΔrtxRID* induced large vesicles, characterizing the protein content of the cholera toxin beta-subunit (CTB).

Consistent with these proteomic differences, we validated these organelles by confocal imaging Caco-2 cells stained with Nile Red and stably expressing Sec61b-miRFP, which respectively represent LDs and ER (Figure 6D-E). The images revealed that WT infection generated migrasomes devoid of ER and LDs signal (figure 6D); In marked contrast, C6706-Δ*rtxRID* infection produced large membrane sheets densely populated with Sec61b-positive puncta, indicating extensive ER entrapment and LDs within these oversized vesicles (Figure 6E). Quantification of lipid droplet content per extracellular structure confirmed a dramatic disparity: whereas WT migrasomes contained barely any detectable lipid droplets, C6706-Δ*rtxRID* derived vesicles harbored an average of 5 to 10 lipid droplets per structure, a difference that was highly statistically significant (Figure 6F). To independently verify the differential enrichment of these lipid droplets associated proteins, we performed immunoblot analysis of purified extracellular vesicles, which confirmed that both PLIN2 and PLIN5 were readily detectable in C6706-Δ*rtxRID* derived vesicles but almost absent from C6706-WT derived migrasomes (Figure 6G). Given that LDs have been reported to exert antimicrobial effects (Bosch et al., 2020), we also observed that *V. cholerae* interacting with LDs appeared elongated (Figure 6H-I), which may represent a sign that the bacterial were being killed. Thus, we searched our proteomics data in aspect of discovery antimicrobial protein which may related to LDs, we found an antimicrobial protein named CHGA is highly enriched in the large vesicles but absent from migrasomes (Figure 6C and 6G).

We also performed immunoblot analysis of purified extracellular vesicles, which confirmed that CXCL2 was readily detectable in C6706-Δ*rtxRID* derived vesicles and relatively less enriched in C6706-WT derived migrasomes (Figure 6G). We next used immunofluorescence microscopy to visualize the spatial distribution of the immunomodulatory cargo within individual vesicles. In Caco-2 cells stably expressing ITGA5-eGFP infected with C6706-Δ*rtxRID*, CXCL2 fluorescence was broadly distributed throughout the large membrane sheets, exhibiting substantially higher integrated intensity and filling a much greater membrane territory. Whereas under C6706-WT infection, the intensity of CXCL2 fluorescence in migrasomes appeared lower (Figure 6J-L). Altogether, we found the unrestricted larger vesicles carry more cargos unfavorable for *V. cholerae*, whereas RID-restricted migrasomes carried less cargos to avoid evoking stronger immune response. Additionally, we take immunoblot assays to detect the cholera toxins delivered from *V. cholerae*, we found that beta subunit of cholera toxin could be detected in both migrasomes and large vesicles (Figure 6M), suggesting that host cells may exploit extracellular vesicles to sequester *V. cholerae* toxin and reduce its cytotoxicity.

### MARTX toxins are required for *V. cholerae* intestinal colonization and virulence gene expression *in vivo*

To determine the contribution of MARTX toxins to *V. cholerae* intestinal colonization in vivo, infant mice were infected with C6706-WT, C6706-Δ*rtxRID*, and C6706-Δ*rtxCPD*. Both mutant strains showed significantly reduced bacterial loads in the ileum compared to wild-type (Figure 7A). Consistent with the colonization defect, transcription of *tcpA* and virulence factor *ctxA* was decreased in both mutant strains (Figure 7B-C). As the RID domain cooperates with the ABH domain to hijack host immune signaling pathways, we next examined the expression of the chemokines in infected ileal tissue. CXCL2 was markedly upregulated (about 6-fold higher) in the Δ*rtxCPD* strain, whereas in the Δ*rtxRID* strain infection, a modest increase (about 2-fold higher) was observed (Figure 7D).

**Figure 7.**
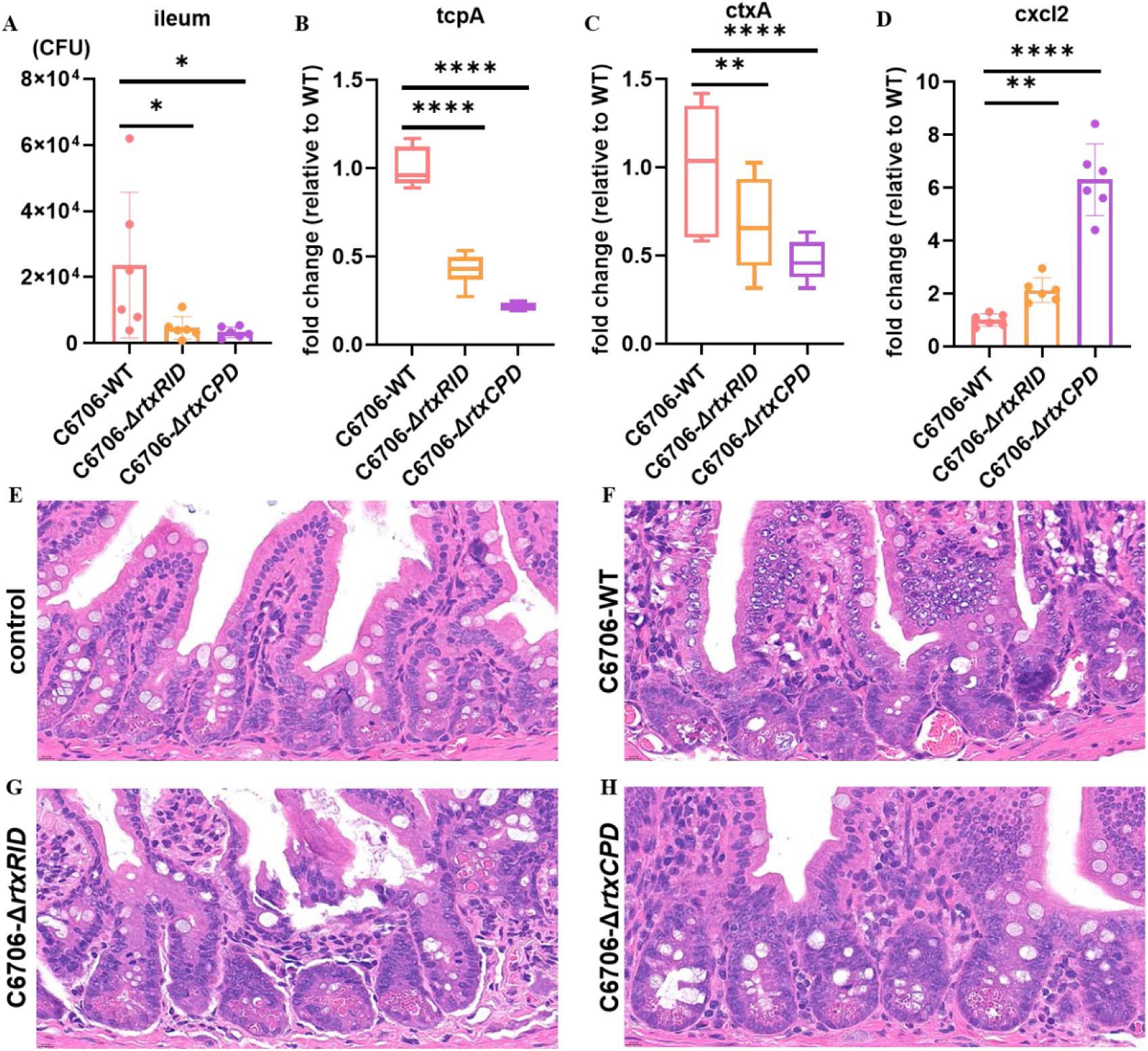
MARTX and the RID effector domain are required for *Vibrio cholerae* colonization and modulate host immune-response in mice. (A) Ileum bacteria load (CFU) in infant mice at 6 hours post infection. Each point represents one animal (n=6 per group). (B) *tcpA*, (C) *ctxA*, and (D) *cxcl2* transcript levels in infant mice ileum measured by qPCR. Each point represents 1 pool of 2 animals. n=6 per pool. P values were calculated using one-way ANOVA followed by Dunnett’s test. (E) - (H) representative H&E-stained sections of mouse ileum at 6 hours post infection: (E) uninfected control, (F) C6706-WT, (G) C6706-Δ*rtxRID*, (H) C6706-Δ*rtxCPD*. Scale bar, 10 μm.

Given the marked differences in bacterial colonization, virulence gene expression, and chemokine production among the three strains, we next examined whether these changes were associated with altered intestinal tissue pathology. Histopathological analysis of H&E-stained ileal sections revealed that mice infected with WT or Δ*rtxRID* strain displayed mucosal damage, including villous blunting, epithelial denudation, and dense inflammatory cell infiltration in the lamina propria, in mice infected with WT (Figure 7F) or Δ*rtxRID* strain (Figure 7G), whereas sections from Δ*rtxCPD*-infected mice were largely indistinguishable from uninfected controls (Figure 7E and 7H). Quantitative scoring showed that WT-infected mice had lower the damage score than Δ*rtxRID*-infected mice, while inflammation scores were only modestly decreased (Figure S7A-B). Δ*rtxCPD*-infected mice showed elevated inflammation relative to uninfected controls, but both inflammation and damage scores were significantly reduced compared with WT-or Δ*rtxRID*-infected mice (Figure S7A-B).

## Discussion

Traditionally, migrasome biogenesis has been regarded as a phenomenon intrinsically coupled to active cell migration. Yet, the field is rapidly expanding beyond autonomous motility, as emerging evidence suggests that non-motile cells can also undergo “non-canonical migracytosis” when challenged by microbial toxins (Li et al., 2024). In this study, we demonstrate that the *V. cholerae* MARTX toxin systematically rewires the host cytoskeleton through the coordinated action of its multiple effector domains, thereby driving migrasome formation in non-migratory intestinal epithelial cells. Deletion of individual effector domains did not merely attenuate host cellular deformation and migrasome production (Figure 2); it also diminished the mutant strains’ adherent capacity to epithelial monolayers and reduced expression of the *tcpA*, an essential determinant of intestinal colonization (Figure 3). These findings imply that MARTX-mediated cytoskeletal reprogramming and migrasome biogenesis are not unidirectional host subjugation events but may instead generate reciprocal fitness benefits for the pathogen, coupling subcellular host manipulation to bacterial colonization potential. Collectively, our findings suggest that the host-pathogen interface during *V. cholerae* infection transcends the simple hijacking of signaling pathways; rather, it constitutes a finely orchestrated subcellular interplay in which migrasome biogenesis and its regulation emerge as a critical nexus for immune modulation and microbial adaptation.

Since its discovery, the ACD domain has been primarily characterized as a driver of cell rounding through covalent cross-linking of G-actin into dimers and multimers (Kudryashov et al., 2008a). Our data reveal an additional, structurally formative role for ACD-mediated actin oligomerization: it serves as the mechanical prerequisite for cell contraction and potentially triggers retraction fibers formation. Long-term confocal live-cell imaging of Lifeact-labeled F-actin further illuminated this process: polymerized actin was transported outward along retraction fibers as cells underwent progressive deformation, ultimately coalescing at distal foci to generate migrasomes (Figure 4B). This observation suggests that MARTX toxins exert consequences not only at the molecular level but also through the subcellular reorganization of host architecture.

While ACD-generated low-order actin oligomers are necessary for retraction formation, they are insufficient for full migrasome maturation. Our data indicate that the ABH domain is required for the subsequent wave of high-order actin polymerization (Figure S3A and Figure 4C). Because ABH functions as a phospholipase unlikely to directly modulate actin dynamics, we hypothesize that it operates indirectly by licensing actin regulatory proteins. Co-immunoprecipitation experiments using either ITGA5 or actin as baits revealed a striking domain-dependent reorganization of the actin interactome: under Δ*rtxABH* infection, polymerized actin associated primarily with FLNA, whereas wild-type infection triggered a wholesale switch toward polymerization-promoting factors including PFN1, DIAPH1, CAP1 and CFL1. ITGA5 retained its association with polymerized actin throughout this remodeling, indicating that integrin-mediated anchorage persists even during extensive cytoskeletal reorganization (Figure 4G). Functional validation through PFN1 depletion marked reduced migrasome formation (Figure S4D-E) and reduced the formation of polymerized actin (Figure 4G), confirming the biological relevance of this remodeled interactome. Through the immunoprecipitation, we can also confirm that ITGA5 only interact with polymerized actin, and transport the polymerized actin outside of the cell eventually forms migrasomes (Figure 4G-H). Combine with literature-guided reasoning suggests that PFN1 activity is normally restrained by phosphatidylinositol 3-phosphate (PI3P) (Saarikangas et al., 2010), and reports that DIAPH1, PFN1 and related factors coordinate actin polymerization(Heisler et al., 2015), we speculate that ABH, in the course of hydrolyzing PI3P to suppress autophagic flux, may concurrently relieve PI3P-mediated inhibition of PFN1 and related regulators, thereby amplifying ACD-driven actin polymerization in a synergistic two-domain mechanism for cytoskeletal remodeling and migrasome maturation. However, this model remains hypothetical, and direct evidence linking ABH-mediated PI3P hydrolysis to PFN1 activation during infection remains to be established.

The structural dynamics of the host cortical actin cytoskeleton are regulated by Rho/Ras family GTPases (Bement et al., 2024), which cycle between active GTP-bound and inactive GDP-bound states to coordinate actin dynamics (Lappalainen et al., 2022). Our data reveal that the RID domain catalyzes the covalent inactivation of multiple Rho/Ras GTPases (Figure 5C), thereby promoting disassembly of the cortical actin meshwork that would otherwise persist during rapid cell traction (Figure S5). This RID-dependent actin remodeling prevents aberrant membrane accumulation and facilitates the formation of discrete migrasomes. Consistent with this model, infection with the Δ*rtxRID strains*, which lack RID-mediated GTPases inactivation, resulted in persistent activation of Rho/Ras GTPases and retention of the cortical actin network. Consequently, cells underwent extensive morphological contraction that produced large, irregular membrane sheets rather than discrete migrasomes. Live-cell imaging further revealed that these sequestered actin structures served as scaffolds stabilizing the irregular membrane patches. Importantly, these large membrane sheets are distinct from migrasomes, which are smaller, more uniform vesicles that form at retraction fibers under normal conditions (Figure S5 and Figure 5A). Moreover, our data suggest H-Ras has a unique role in converting these ruptured, irregular membrane patches into spherical large vesicles, potentially allowing them to be released from the cell in a more stable conformation. This conversion appears specific to H-Ras, as the other GTPases tested did not similarly promote formation of spherical large vesicles (Figure 5D).

Beyond their established roles in cell-cell communication and embryonic development, migrasomes are emerging as critical conduits for immune regulation. In our *V. cholerae* infection model, the RID domain functions as a morphological and compositional sieve that governs the quality and quantity of extracellular release. Critically, while both wild-type migrasomes and C6706-Δ*rtxRID* induced blebs contain the pro-inflammatory chemokine CXCL2, the latter sequester substantially higher amounts and exhibit broader spatial distribution throughout the expanded membrane territory. We propose that this domain-dependent cargo restriction severs a dual purpose for the pathogen: by constraining vesicle dimensions and excluding bulky organelles, RID simultaneously attenuates the release of antimicrobial proteins, endogenous danger signals and limits the inflammatory broadcast volume, thereby dampening early immune surveillance to facilitate bacterial colonization (Figure 6).

The potential mechanistic link between RID-dependent cargo restriction and virulence is further supported by divergent survival phenotypes in animal infection models. Previous studies demonstrated that Δ*rtxRID* mutant infection leads to increased host survival, whereas Δ*rtxABH* infection accelerates lethality (Gavin and Satchell, 2019). In our infant mouse infection model, defects in RID or overall MARTX function were associated with reduced bacterial colonization and decreased *tcpA* transcription (Figure 7A-B), correlating with our in vitro infection assays (Figure 3A-D). Elevated *cxcl2* transcription in infected ileal tissue (Figure 7D) was also consistent with our immunofluorescence and Western blot analyses (Figure 6J-L). Because both wild-type and Δ*rtxRID* strains retained the capacity to induce rapid cell traction, histopathological analysis revealed substantial tissue damage in both groups, with only modestly reduced inflammation and damage scores in WT-infected mice compared with Δ*rtxRID*-infected mice (Figure 7F-G and Figure S7). These observations prompt us to hypothesize that the distinct migrasomes and large vesicles phenotypes observed in vitro may contribute to the differential inflammatory and colonization outcomes seen in vivo. However, our current data do not establish a causal link between migrasome biogenesis and the *in vivo* phenotypes; whether *V. cholerae*-induced migrasomes directly activate resident macrophages, serve as chemotactic beacons for immune cell recruitment, or function in antigen presentation during later infection stages remains an important open question. Further studies utilizing genetic models deficient in migrasome formation, such as *tspan4-*or *tspan9*-deficient mice, will be essential to determine the necessity of migrasome biogenesis for host defense against *Vibrio* and to profile the dynamics of intestinal immune cell recruitment across MARTX domain variants. Collectively, our findings reveal that the host-pathogen interface is defined by a mechanical tug-of-war over migrasome biogenesis: the pathogen deploys RID to restrict and attenuate extracellular inflammatory signals, while the host maintains the capacity to transport actin and chemokines along retraction fibers, establishing the foundation for eventual pathogen clearance.

## Methods and Materials

### Cells, Strains and Plasmids

Cells, strains and plasmid constructs are listed in Table S1, Table S2, and Table S3 respectively. To construct deletion mutants of *V. cholerae*, allelic exchange was carried out using the counter selectable suicide plasmids pWM91 and pDS132. For each target gene, two ∼800-bp homology arms flanking the 5′ and 3′ ends of the coding sequence were PCR-amplified and fused into a linearized suicide vector by Gibson assembly. The resulting plasmids were transferred into *V. cholerae*, and double-crossover recombinants were recovered on LB agar supplemented with 10% sucrose. To construct stable cell line, we packaged lentivirus in 293T by transfecting plasmids pLenti, psPAX and pMD2.G in a ratio of 4:3:1 for 48 hours. The lentivirus was collected and used to infect cells for 48 hours. Cells with successfully integrated genes were selected under puromycin.

### Antibodies, Reagents, and Growth Conditions

For the staining assay, we performed WGA (Servicebio) for membrane labeling, Nile Red (Beyotime) for lipid droplets and neutral fatty acid labeling, DAPI (Beyotime) and Hoechst (Beyotime) for nuclear labeling, polybrene (Merck millipore) for promoting lentivirus infection efficiency, puromycin (Sigma-Aldrich) for stable cell line selection, and SYBR (Vazyme) for RT-qPCR. For Western blot, we utilize β-actin (Abbkine), vinculin (HUABIO), ITGA5 (HUABIO), RpoB (BioLegend), PFN1 (HUABIO), DIAPH1 (HUABIO), CFL1 (HUABIO), CAP1 (HUABIO), FLNA (HUABIO), CXCL2 (CST), PLIN5 (HUABIO), PLIN2 (HUABIO), HA (HUABIO), Flag (HUABIO) as primary antibodies and HRP conjugated mouse or rabbit (proteintech) as secondary antibodies. For the immunofluorescence assay, CXCL2 was used as the primary antibody and Dylight 594, Goat Anti-Rabbit IgG (Abbkine) as secondary antibodies.

Bacteria strains were routinely cultured in LB medium or on LB agar with appropriate antibiotics at 37°C except *Vibrio alginolyticus* and *Vibrio harveyi* grow at 30°C. Cells were routinely cultured in DMEM (VivaCell) supplemented with 10% FBS (WISENT) and 1% Penicillin-streptomycin dual antibiotics (Gibco).

### Bacterial infection of cultured cells

Unless otherwise specified, Caco-2 cells were infected with bacteria at a multiplicity of infection (MOI) of 10 (bacteria: cell) for the indicated times. Bacteria grown to mid-log phase (OD600 ≈ 0.4) in LB medium were diluted in PBS, resuspended in DMEM without antibiotics, and added to cells. Infected cells were incubated at 37°C in a 5% CO2 humidified incubator. After infection, cells were washed twice with PBS and processed for microscopy, Western blotting, RNA extraction, or migrasome purification as described below.

### Dual sgRNA CRISPR-Cas9 Knockout Assay

sgRNAs targeting ITGA5 were designed using CRISPor.com and cloned into the fragment which was arranged as “sgRNA scaffold – U6 promoter” with primers: F-5’ aaacgtctctcaccgTCATGGATCGGACCCCTGACgttttagagctagaaatagcaagttaaaa - 3’ and R-5’ CGTCTCCaaacAGACCCTGCTCATCCAGAATCCGGTGTTTCGTCCTTTCCAC-3’. The fragment and pLentiCRISPRv2 were cut by BsmBI (NEB). The fragment was inserted into the plasmid by using T4 ligase (Takara). Constructed plasmids were packaged into lentivirus and integrated into Caco-2 genome. A single cell clone was collected and cultured until determined by Western blot and PCR.

### Recombinant Protein Preparation

Full-length ACD domain (residues S1972–L2447) from *V. cholerae* MARTX (MARTXvc), fused with an N-terminal Flag tag, was cloned into the pET-28a vector. The recombinant plasmid was transformed into Escherichia coli BL21 (DE3) cells, which were cultured in LB medium supplemented with 50 μg/mL kanamycin at 37°C until the OD600 reached 0.8.

Protein expression was induced by adding 0.1 mM IPTG, followed by incubation at 20°C for 16 h. Bacterial cells were harvested by centrifugation at 5,000 × g for 15 min and resuspended in lysis buffer (25 mM Tris-HCl, 150 mM NaCl, pH 7.6). Cells were lysed using a high-pressure homogenizer, and the lysate was clarified by centrifugation at 12,000 × g for 30 min. The soluble fraction was loaded onto Ni-NTA agarose resin (BBI) and washed with lysis buffer containing 50 mM imidazole. Bound proteins were eluted with lysis buffer containing 300 mM imidazole.

Human β-actin (*ACTB*) with an N-terminal Strep-tag II was cloned into the pcDNA3.1 vector and transiently transfected into 293F cells using PEI. Transfected cells were cultured in DMEM supplemented with 10% FBS at 37°C in a 5% CO2 humidified incubator for 24 h. Cells were harvested by centrifugation at 1,200 × g for 15 min, washed once with PBS, and resuspended in buffer A (20 mM Tricine, 150 mM NaCl, 1 mM EDTA, 1 mM EGTA, protease inhibitor cocktail, pH 7.8). Cells were lysed by sonication on ice, and the lysate was clarified by centrifugation at 12,000 × g for 30 min to remove cell debris and membranes. The cleared supernatant was loaded onto Strep-Tactin Sepharose resin (AlpaLifeBio) and washed with buffer A. Bound Strep-tagged ACTB was eluted with buffer (25 mM Tris-HCl, 150 mM NaCl, pH 7.6) supplemented with 2.5 mM desthiobiotin.

For both proteins, affinity-purified eluates were concentrated and further purified by size-exclusion chromatography on a Superdex™ 75 Increase 10/300 GL column (Cytiva) equilibrated with SEC buffer (25 mM Tris, 150 mM NaCl, pH 7.6).

### Animal ethics statement

Experiments involving mouse materials were approved by the Institutional Animal Care and Use Committees of Nanjing Agricultural University. The Nanjing Agricultural University animal management program is accredited by the Laboratory Animal Monitoring Committee of Jiangsu, Province, China.

### Infant Mouse Model

CD-1 3∼5-day old infant mice (Charles River) were purchased from Vital River company were separated from their dams for 4 hours prior to oral inoculation with 10^8^ CFU bacteria in 50 μl LB. After *V. cholerae* gavage, mice were sacrificed at 6 hours and ileum was separated for RNA extraction, CFU and histopathological analysis.

### In vitro Cross-link Assay

Purified strep-ACTB and 6His-RID-flag was mixed at ratio 100:1 supplemented with 1 mM MgCl2 and 250 μM ATP. Reaction was performed at 37°C for 30 minutes. The reaction was ended by adding SDS-loading and boiled at 95°C for 10 minutes.

### Negative-stain Transmission Electron Microscopy

10 μL purified migrasomes or large vesicles (0.1 mg/mL) were incubated with glow discharged Quantifoil R 2/2 Cobe 200 mesh grid for 10 minutes followed by staining uranium acetate 10 minutes. Samples were subject into 120 kv transmission electron microscopy (thermos fisher) and data collection at 53,000 × magnifications.

### Confocal Microscopy

cells were seeded on confocal chamber (NEST) and transferred to a Ziess LSM900 confocal microscope equipped with a live-cell imaging system. Images were aquired using 63 × objective at 1980 × 1980 pixels in Airyscan model. For long-term time-lapse experiments, cells were infected with *V. cholerae* and image immediately. Images were aquired at 3 min intervals for indicated time periods.

### Migrasomes and Large Vesicles Purification

The purification of migrasomes or lager vesicles mainly refers (Jiang et al., 2023). In brief, Caco-2 cells were cultured at 25 cm dishes and infected with either C6706-WT or C6706-Δ*rtxRID* once they reach ∼60% confluence. After 3 to 4 hours of infection, the cells were removed the cell culture and wash twice by PBS (Vivacell) then digested with trypsin supplemented with 0.25% EDTA (Gibco) for 4 minutes. The digestion was ended by adding DMEM supplemented with 10% FBS. Cells were collected and centrifuge at 800 rpm to remove eukaryotic cells and 4000 rpm to remove *V. cholerae*. Then the supernatant was subject to ultracentrifugation by using SW-41 Ti (Beckman) at 20,000 × g. The pellets were collected and subject to an iodixanol (BasalMedia) gradient (5%, 10%, 15%, 20%, 25%, 30%) followed by ultracentrifugation at 150,000 × g using SW-41 Ti.

### Click Chemistry

Click chemistry mainly referred to (Zhou et al., 2017). In brief, to detect the fatty acylation of Rho/Ras GTPases by RID in cells, Caco-2 cells stably expressing the indicated GTPases were generated by lentiviral transduction of the corresponding GTPase-encoding plasmids. The cells were subsequently infected with C6706-WT or C6706-Δ*rtxRID* at an MOI of 10. At 30 min post-infection, the culture medium was supplemented with 30 μM alk-16. Following 4 h of metabolic labeling, the cells were harvested and lysed in lysis buffer (25 mM HEPES, pH 7.6, 150 mM NaCl, and 1% Triton X-100). HA-tagged Rho/Ras GTPases were immunoprecipitated from the lysates using anti-HA antibody-coated beads at 4°C. After washing three times with wash buffer (25 mM HEPES, pH 7.6, 150 mM NaCl), the proteins were subjected to click chemistry in 20 μl reaction buffer (25 mM HEPES, pH 7.6, 150 mM NaCl, and 4% SDS) containing 200 μM azido-biotin, 200 μM Tris[(1-benzyl-1H-1,2,3-triazol-4-yl)methyl]amine, 2 mM CuSO4, and 2 mM TCEP at 30°C for 3 h in the dark. Chloroform/methanol precipitation was performed to remove excess azido-biotin. The protein pellets were washed three times with methanol, air-dried, and resuspended in SDS sample loading buffer. The samples were resolved by 12% SDS-PAGE and analyzed by Western blotting using a streptavidin-biotinylated HRP complex (MCE).

### Immunoprecipitation Assays

To examine whether polymerized actin interacts with ITGA5 in vivo, the infected cells were washed with PBS and lysed with lysis buffer that contained 25 mM HEPES, pH 7.6, 150 mM NaCl, and 1% Triton-X100. After centrifugation at 12000 rpm for 10 min, the supernatant of the cell lysate was collected and mixed with beads which cultured with HA antibody. After incubation at 4°C overnight, the beads were washed five times with lysis buffer and denatured in SDS sample loading buffer. The samples were subjected to SDS-PAGE followed by a Western blot assay: 8% SDS-PAGE to identify polymerized actin, 10% SDS-PAGE to identify ITGA5, FLNA, DIAPH1, CAP1, and vinculin, 12% SDS-PAGE to identify PFN1 and CFL1.

### RNA Extraction

In the bacteria adhesion assay, Caco-2 cells were cultured on 24-well dishes and infected with various genotype *V. cholerae* once they reached ∼60% confluence. After 2 or 3 hours of infection, cells together with *V. cholerae* were dissolved in trizol (Invivogen). In vivo infection assay, infant mouse ileum was collected after various genotype *V. cholerae* infection. The tissue was soaked in 1 ml trizol and subjected to a tissue homogenizer for 1.5 minutes grinding three times with 0.5-minute intervals. Then chloroform-isopropanol was used to RNA precipitate. The RNA precipitate was washed twice with 75% ethanol and air-dried then resolved in DEPC water.

### Quantitative real-time PCR (qPCR)

Reverse transcription was performed using HiScript III RT SuperMix (Vazyme) according to the manufacturer’s instructions. The resulting cDNA was diluted 5-fold and used as the template for qPCR. Real-time qPCR was performed on a LightCycler 96 System (Roche). All reactions were run in triplicate. Relative expression levels were normalized to internal control genes: *V. cholerae 16S* rRNA, Caco-2 *18S* rRNA, or mouse *gapdh* and calculated using the 2^-ΔΔCt method.

### Immunofluorescence assay

Cell glass slides (NEST) were placed on a 24-well cell culture dish and coated with 1% extracellular matrigel (Corning) for two hours before seeding Caco-2 cells. The cells were infected with either C6706-WT or C6706-Δ*rtxRID* until they reached ∼60% confluence. Cells were fixed with polyformaldehyde for 20-30 minutes. The cells membrane was mildly dissolved under incubation of 0.05% saponin for 15-20 minutes, 5% BSA was used as block regent. Both primary antibodies and secondary antibodies were incubated for 1 hour. After air-drying, the cell glass slides were inverted onto the slides with fix regent supplemented with DAPI.

### Bacteria adhesion assay

Caco-2 cells were seeded on a 24-well cell dish and infected with various genotype *V. cholerae* until they reach ∼60% confluence. After 2-or 3-hour infection, the cell culture was removed and the cells were washed with PBS two times to exclude the free-floating bacteria. The cells were digested form the dish by incubating trypsin supplemented with EDTA for 3 minutes and ended by adding DMEM supplemented with 10% FBS. Cells were collected into a centrifuge tube with a final volume of 1 mL, followed by gradient dilution in PBS. The amount of bacterial adhesion to cells was measured as CFU.

### Cryo-ET Samples Preparation

Caco-2 cells were infected with *V. cholerae* until they reached ∼40-50% confluence on EM grids (Quantifoil Au R2/1, 200 meshes). After 3-4 hours of infection, cells were inspected under the light microscope to ensure they had successfully grown on the EM grids. Then, 3.5 μl of concentrated (10×) gold tracers (Ted Pella, USA) severing as fiducial markers were applied to the gold grids before they were manually blotted twice and then plunge-frozen into liquid ethane using a Vitrobot Mark IV (Thermo Fisher Scientific, USA) at liquid nitrogen temperature (Tivol et al., 2008). Grids were stored in liquid nitrogen before they were transferred to the columns of the cryo-electron microscopes for imaging.

*Vibrio cholerae* was incubated with migrasomes purified from infected Caco-2 cells at 37°C for 2 hours. 9 μl of the incubation system was mixed with 1 μl concentrated (10×) gold tracers, which were applied to the EM grids (Quantifoil Cu R2/2, 300 meshes) before they were manually blotted twice and then plunge-frozen into liquid ethane using a Vitrobot Mark IV as mentioned above.

### Cryo-ET data collection

Tilt series of *V. cholerae* infected Caco-2 cells and *V. cholerae* incubated with purified migrasomes were acquired in a 300 kV Titan Krios microscope (Thermo Fisher Scientific, USA) at a magnification of 26,000× on a Falcon IV camera (Thermo Fisher Scientific, USA). Only cells with migrasome regions within the grid squares were picked for imaging. For each chosen cell, five to ten tilt series were collected on randomly selected cellular areas. Tilt series were acquired at a 3° interval from −60° to +60° using a dose-symmetric tilting scheme in SerialEM (Hagen et al., 2017; Mastronarde, 2005) with a 15eV slit inserted. The cumulative dose per tilt series was ∼120 e^−^/Å^2^. The defocus was set from −8.0 to −10.0 μm. The pixel size was 3.343 Å/pix for the Falcon IV camera.

### Tomographic reconstruction and denoising

Before reconstruction, the quality of all acquired tilt series was manually inspected, and data suffering from bad stage drift were discarded. After that, the remaining tilt series were aligned with either the fiducial-based or the patch-tracking method implemented in the IMOD package (Kremer et al., 1996). The tomographic reconstruction was performed using either the weighted back projection (WBP) method or the simultaneous iterative reconstruction technique (SIRT) implemented in IMOD (Agulleiro and Fernandez, 2011), yielding 3D reconstructed subcellular volumes called tomograms (4k × 4k × 1k). To enhance the contrast, the reconstructed tomograms were binned by two in EMAN2 (Tang et al., 2007) followed by noise reduction using the nonlinear anisotropic diffusion (NAD) tool in IMOD (Frangakis and Hegerl, 2001).

### Tomogram Visualization and Segmentation

We used Amira software (Thermo Fisher Scientific, USA) to visualize and annotate migrasome-containing tomograms after reconstruction and filtering. Membranous structures such as migrasomal membranes, bacterial membranes, secreted blebs, Endoplasmic Reticulum (ER), Golgi Complex membranes, etc, were segmented with thresholding and brush tools in Amira. The interpolation method was used to facilitate segmentation. A membrane enhancement filter was applied to all segmented membranous structures. Filamentous structures like retraction fibers were either segmented with EMAN2, followed by refinement in Amira, or segmented manually in Amira. Other components, such as migrasomal content, bacterial cytoplasm, and protein-like densities connecting bacteria and migrasomes, were manually segmented in Amira with sufficient care.

### Mass spectrometry analysis

Migrasomes or large vesicles and actin immunoprecipitation samples were sent to Chi Biotech (Shenzhen) Co Ltd. for mass spectrometry analysis.

Migrasomes and large vesicles were sent for DIA-based quantitative proteomic analysis. Protein extraction, denaturation, reduction, alkylation, tryptic digestion, and peptide cleanup were performed using the iST Sample Preparation kit (PreOmics) according to the manufacturer’s protocol. The resulting peptides were reconstituted in 0.1% formic acid and separated by UHPLC on a C18 column using an acetonitrile gradient. Eluted peptides were analyzed on a timsTOF HT mass spectrometer (Bruker Daltonik) in diaPASEF mode. Raw DIA data were processed with Spectronaut 19 (Biognosys) against the UniProt *Homo sapiens* database (2025 release, 20,656 entries). Trypsin was specified as the digestion enzyme, carbamidomethylation of cysteine as the fixed modification, and oxidation of methionine and N-terminal acetylation as variable modifications. Protein quantities were calculated using the MaxLFQ algorithm with local normalization. Precursor, peptide, and protein identifications were filtered at a 1% false discovery rate.

Actin immunoprecipitated from uninfected and C6706-WT-infected Caco-2 cells was sent for proteomic analysis to a commercial service provider. Immunoprecipitated proteins were separated by SDS-PAGE, and gel bands were cut into small pieces, destained, reduced with dithiothreitol, alkylated with iodoacetamide, and digested overnight with trypsin. The resulting peptides were extracted from the gel pieces, desalted using Pierce C18 spin tips, and dried in a SpeedVac concentrator. The peptides were reconstituted in 0.1% formic acid and analyzed by an EASY-nanoLC 1200 system coupled to an Orbitrap Exploris 480 mass spectrometer (Thermo Fisher Scientific) operating in data-dependent acquisition (DDA) mode. Tandem mass spectra were processed with PEAKS Studio 13 (Bioinformatics Solutions Inc.) against the UniProt *Homo sapiens* database (2025 release, 20,663 entries). Trypsin was specified as the digestion enzyme, carbamidomethylation of cysteine as the fixed modification, and oxidation of methionine, deamidation of asparagine and glutamine, and N-terminal acetylation as variable modifications. Peptide and protein identifications were filtered at a 1% FDR, and proteins were required to contain at least one unique peptide.

### Histological analysis (H&E staining)

Mouse small intestine samples were collected and fixed in 4% paraformaldehyde (PFA) at room temperature for 24 h and sent to Servicebio (Wuhan) Co Ltd. for H&E staining. In Brief, fixed tissues were washed with PBS, dehydrated through a graded ethanol series, cleared in xylene, and embedded in paraffin wax. Four-micrometer-thick sections were prepared using a rotary microtome, mounted on glass slides, and baked at 60°C for 30 min. The sections were dewaxed in xylene, rehydrated through descending concentrations of ethanol, and rinsed in distilled water. The sections were stained with hematoxylin for 3–5 min, differentiated in 1% acid alcohol, blued in running tap water, and counterstained with eosin for 1–3 min. After dehydration through ascending ethanol and clearing in xylene, the slides were mounted with neutral balsam and visualized under a light microscope.

### Histopathological scoring

H&E-stained ileal sections were evaluated independently by three investigators in a blinded manner. Two parameters were assessed for each section: inflammation and tissue damage, both scored on a scale of 0 to 5. For the inflammation score, five features were evaluated independently: neutrophil infiltration, lymphocyte infiltration, macrophage infiltration, eosinophil infiltration, and lamina propria inflammatory expansion. Each feature was scored from 0 (absent) to 5 (severe/diffuse). The final inflammation score for each section was calculated as the mean of the five individual sub-scores. Tissue damage was assessed as a single composite parameter based on overall architectural disruption, including villous blunting, epithelial denudation, crypt distortion, submucosal edema, and mucosal ulceration/hemorrhage, and scored from 0 (absent) to 5 (severe/diffuse).

### Statistical analysis

Data are presented as mean ± standard error of the mean (SEM), with the exact sample size (n) for each experiment provided in the corresponding figure legends. For comparisons involving three or more groups, statistical significance was determined by one-way analysis of variance (ANOVA) followed by Dunnett’s multiple comparisons test. For two-group comparisons, an unpaired two-tailed Student’s t-test was used. Densitometric quantification of Western blot bands, fluorescence intensity line scans, and manual counting of vesicles or lipid droplets were performed using ImageJ (National Institutes of Health). qPCR data and bacterial CFU counts were log-transformed when appropriate before statistical analysis. Histopathological scoring of H&E-stained ileal sections was performed independently by three investigators in a blinded manner, and the composite inflammation and epithelial damage scores were analyzed by one-way ANOVA followed by Dunnett’s test. All statistical analyses were conducted using GraphPad Prism 8 (GraphPad Software). A P-value < 0.05 was considered statistically significant. No data points or animals were excluded from the analysis unless pre-established exclusion criteria were applied

## Acknowledgments

This work was supported by National Natural Science Foundation of China (32270061 and 32300597). Yang Fu is an investigator of SUSTech Institute for Biological Electron Microscopy. We would like to thank the cryo-EM center of Southern University of Science and Technology for helping us with our protein structure analysis.

## Author Contributions

conceptualization PY. X. and Y. F.; methodology PY. X. and Y. F.; investigation, all authors; writing-original draft and review and editing, PY. X., M. L., R. W., M. T. and Y. F.

