## Supplementary Information for "A host-pathogen tug-of-war over migrasome biogenesis in *Vibrio cholerae* infection"

**Table S1 Strains used in this study. Related to Methods “Bacterial strains”.**

| organisms/strains | genotype | description | source |
|---|---|---|---|
| *Vibrio cholerae* C6706 | parental | O1 toxigenic strain with resistance to streptomycin or streptomycin /rifampicin | Lab stock |
| | Δ*ctxAB* | cholera toxin subunit A and B knock out used for screening toxins trigger cell migracytosis | This study |
| | Δ*hly/hapA* | hemolysin and hapA knock out used for screening toxins trigger cell migracytosis | Lab stock |
| | Δ*vipA* | Type 6 secretion system structure gene *vipA* knock out used for screening toxins trigger cell migracytosis | This study |
| | Δ*rtxCPD* | *MARTX* CPD domain knocks out for screening toxins trigger cell migracytosis and in vitro or in vivo infection assay | This study |
| | Δ*rtxABH* | *MARTX* ABH domain knocks out for cell migracytosis researching and in vitro infeciton assay | This study |
| | Δ*rtxACD* | *MARTX* ACD domain knocks out for cell migracytosis researching and in vitro infeciton assay | This study |
| | Δ*rtxRID* | *MARTX* RID domain knocks out for cell migracytosis researching and in vitro or in vivo infection assay | This study |
| *Vibrio vulnificus* 55 | parental | Streptomycin-resistant strain | Lab stock |
| *Vibrio furnissii* VFBJ02 | parental | Ampicillin resistant strain | Lab stock |
| *Vibrio harveyi* BB170 | parental | A kanamycin resistant strain, culture at 30℃ | Lab stock |
| *Vibrio parahaemolyticus* 1360 | parental | Streptomycin resistant strain | Lab stock |
| *Vibrio alginolyticus* SH9-90 | parental | Rifampin-resistant strain, culture at 30℃ | Lab stock |
| *Vibrio mimicus* | parental | Rifampin-resistant strain | Lab stock |
| *E. coli* BL21(DE3) | F- dcm ompT hsdS gal λ | Host bacteria used for protein purification | Lab stock |
| *E. coli* DH5α | K12-derivative laboratory strain | Host bacteria used for constructing recombination plasmids | Lab stock |
| *E. coli* BW20676λ*pir* | K12-derivative laboratory strain | For the construction of gene-knockout vector and mating assays | Lab stock |
| *E. coli* MG1655 | Parental | Streptomycin-resistant strain for infection assay | Lab stock |

**Table S2 Cells used in this study. Related to Methods "Mammalian Cells".**

| organisms/cells | genotype | description | source |
|---|---|---|---|
| Caco-2 | parental | The human clonal colonic adenocarcinoma cell line, structural and functional characteristics resemble those of small intestinal epithelial cells | lab store |
| | *ITGA5*$^{-/-}$ | integrin alpha 5 gene knock out, used for identify migrasomes | This study |
| | TSPAN4-mCherry | stably express TSPAN4 recombinant with mCherry, used as bio-marker for migrasomes | This study |
| | ITGA5-eGFP | stably express ITGA5 recombinant with eGFP, used as bio-marker for migrasomes | This study |
| | ITGA5-eGFP/ lifeact-miRFP | stably express ITGA5 recombinant with eGFP and lifeact recombinant with miRFP, used as bio-marker for migrasomes and polymerized actin | This study |
| | *ITGA5*$^{-/-}$/ lifeact-miRFP | integrin alpha 5 gene knock out and stably express lifeact recombinant with miRFP, used for track polymerized actin in migracytosis defective cells | This study |
| | shPFN1 | profilin 1 gene knock down, used for actin polymerization assay | This study |
| | shFLNA | filamin-A gene knock down, used for actin polymerization assay | This study |
| | shCAP1 | Adenylyl cyclase-associated protein 1 gene knock down, used for actin polymerization assay | This study |
| | shDIAPH1 | Protein diaphanous homolog 1 gene knock down, used for actin polymerization assay | This study |
| | shVASP | Vasodilator-stimulated phosphoprotein gene knock down, used for actin polymerization assay | This study |
| | shPFN1/ TSPAN4-mCherry/ lifeact-miRFP | profilin 1 gene knock down, stably express TSPAN4-mCherry and lifeact-miRFP, used for tracking polymerized actin in PFN1 knock down cells | This study |
| | ITGA5-eGFP/ mCherry-H-Ras | stably express ITGA5 recombinant with eGFP and H-Ras recombinant with mCherry, used for vesicle morphology observation assay | This study |
| | ITGA5-eGFP/ mCherry-RhoC | stably express ITGA5 recombinant with eGFP and RhoC recombinant with mCherry, used for vesicle morphology observation assay | This study |
| | ITGA5-eGFP/ lifeact-miRFP/ | stably express ITGA5 recombinant with eGFP and Rac3 recombinant with mCherry, used for | This study |

| | | | |
|---|---|---|---|
| | mCherry-Rac3 | vesicle morphology observation assay | |
| | ITGA5-eGFP/ lifeact-miRFP/ mCherry-RhoG | stably express ITGA5 recombinant with eGFP and RhoG recombinant with mCherry, used for vesicle morphology observation assay | This study |
| | HA-RhoB | stably express RhoB with HA tag, used for click chemistry | This study |
| | HA-RhoC | stably express RhoC with HA tag, used for click chemistry | This study |
| | HA-Rac1 | stably express Rac1 with HA tag, used for click chemistry | This study |
| | HA-Rac3 | stably express Rac3 with HA tag, used for click chemistry | This study |
| | HA-H-Ras | stably express H-Ras with HA tag, used for click chemistry | This study |
| | HA-R-Ras | stably express R-Ras with HA tag, used for click chemistry | This study |
| | HA-R-Ras2 | stably express R-Ras2 with HA tag, used for click chemistry | This study |
| | HA-Rap2A | stably express Rap2A with HA tag, used for click chemistry | This study |
| | HA-Rap2B | stably express Rap2B with HA tag, used for click chemistry | This study |
| | Sec61b-miRFP | stably express Sec61b recombinant with miRFP, used for labeling ER | This study |
| A549 | ITGA5-eGFP | The human lung adenocarcinoma cell line, stably express ITGA5 recombinant with eGFP | This study |
| U2OS | ITGA5-eGFP | The human osteosarcoma cell line, stably express ITGA5 recombinant with eGFP | This study |
| 293T | ITGA5-eGFP | The human embryonic kidney cell line, stably express ITGA5 recombinant with eGFP | This study |
| L929 | ITGA5-eGFP | The mouse fibroblast cell line, stably express ITGA5 recombinant with eGFP | This study |

**Table S3 Plasmids used in this study. Related to Methods “plasmids”.**

| Name | Description | Usage | Source |
|---|---|---|---|
| pDS132 | *oriV*$_{R6K}$-containing suicide vector, Cm$^R$ | Backbone plasmid | lab store |
| pDS132-Δ*ctxAB* | pDS132 derived plasmid containing 0.8 kb upstream of *ctxA* and 0.8 kb downstream of *ctxB* in its MCS | Used to delete *ctxAB* from C6706 | This study |
| pDS132-Δ*vipA* | pDS132 derived plasmid containing 0.8 kb upstream and downstream of *vipA* in its MCS | Used to delete *vipA* from C6706 | This study |
| pDS132-Δ*rtxCPD* | pDS132 derived plasmid containing 0.8 kb upstream and downstream of CPD domain in *MARTX* in its MCS | Used to delete CPD domain in *MARTX* from C6706 | This study |
| pDS132-Δ*rtxABH* | pDS132 derived plasmid containing 0.8 kb upstream and downstream of ABH domain in *MARTX* in its MCS | Used to delete ABH domain in *MARTX* from C6706 | This study |
| pDS132-Δ*rtxACD* | pDS132 derived plasmid containing 0.8 kb upstream and downstream of ACD domain in *MARTX* in its MCS | Used to delete ACD domain in *MARTX* from C6706 | This study |
| pDS132-Δ*rtxRID* | pDS132 derived plasmid containing 0.8 kb upstream and downstream of RID domain in *MARTX* in its MCS | Used to delete RID domain in *MARTX* from C6706 | This study |
| psPAX2 | a second-generation lentiviral packaging plasmid that expresses the HIV-1 gag and pol genes | Used to package pLenti plasmids | lab store |
| pMD2.G | a lentiviral envelope plasmid encoding the vesicular stomatitis virus glycoprotein (VSV-G). | Used to package pLenti plasmids | lab store |
| pLenti | a lentiviral transfer plasmid designed for stable integration and long-term expression of target genes | Backbone plasmid | lab store |
| pLenti-ITGA5-eGFP | pLenti derived plasmid containing integrin alpha 5 gene cloned from Caco-2 cDNA library fused with eGFP | used to construct stably express ITGA5-eGFP cell line in Caco-2 | This study |
| pLenti-TSPAN4-mCherry | pLenti derived plasmid containing Tetraspanin-4 gene cloned from Caco-2 cDNA library fused with mCherry | used to construct stably express TSPAN4-mCherry cell line in Caco-2 | This study |
| pLenti-lifeact-miRFP | pLenti derived plasmid containing synthesized lifeact gene fused with miRFP | used to construct stably express lifeact-miRFP cell line in Caco-2 | This study |
| pLenti-mCherry-H-Ras | pLenti derived plasmid containing mCherry fused with H-Ras gene cloned from Caco-2 | used to construct stably express | This study |

| | | | |
|---|---|---|---|
| | cDNA library | mCherry-H-Ras cell line in Caco-2 | |
| pLenti-mCherry-RhoC | pLenti derived plasmid containing mCherry fused with RhoC gene cloned from Caco-2 cDNA library | used to construct stably express mCherry-RhoC cell line in Caco-2 | This study |
| pLenti-mCherry-RhoG | pLenti derived plasmid containing mCherry fused with RhoG gene cloned from Caco-2 cDNA library | used to construct stably express mCherry-RhoG cell line in Caco-2 | This study |
| pLenti-mCherry-Rac3 | pLenti derived plasmid containing mCherry fused with Rac3 gene cloned from Caco-2 cDNA library | used to construct stably express mCherry-Rac3 cell line in Caco-2 | This study |
| pLenti-HA-RhoB | pLenti derived plasmid containing HA-tag fused with RhoB gene cloned from Caco-2 cDNA library | used to construct stably express HA-RhoB cell line in Caco-2 | This study |
| pLenti-HA-RhoC | pLenti derived plasmid containing HA-tag fused with RhoC gene cloned from Caco-2 cDNA library | used to construct stably express HA-RhoC cell line in Caco-2 | This study |
| pLenti-HA-Rac1 | pLenti derived plasmid containing HA-tag fused with Rac1 gene cloned from Caco-2 cDNA library | used to construct stably express HA-Rac1 cell line in Caco-2 | This study |
| pLenti-HA-Rac3 | pLenti derived plasmid containing HA-tag fused with Rac3 gene cloned from Caco-2 cDNA library | used to construct stably express HA-Rac3 cell line in Caco-2 | This study |
| pLenti-HA-H-Ras | pLenti derived plasmid containing HA-tag fused with H-Ras gene cloned from Caco-2 cDNA library | used to construct stably express HA-H-Ras cell line in Caco-2 | This study |
| pLenti-HA-R-Ras | pLenti derived plasmid containing HA-tag fused with R-Ras gene cloned from Caco-2 cDNA library | used to construct stably express HA-R-Ras cell line in Caco-2 | This study |
| pLenti-HA-R-Ras2 | pLenti derived plasmid containing HA-tag fused with R-Ras2 gene cloned from Caco-2 cDNA library | used to construct stably express HA-R-Ras2 cell line in Caco-2 | This study |
| pLenti-HA-Rap2A | pLenti derived plasmid containing HA-tag fused with Rap2A gene cloned from Caco-2 | used to construct stably express HA- | This study |

| | | | |
|---|---|---|---|
| | cDNA library | Rap2A cell line in Caco-2 | |
| pLenti-HA-Rap2B | pLenti derived plasmid containing HA-tag fused with Rap2B gene cloned from Caco-2 cDNA library | used to construct stably express HA-Rap2B cell line in Caco-2 | This study |
| pLenti-Sec61b-miRFP | pLenti derived plasmid containing Protein transport protein Sec61 subunit beta gene cloned from Caco-2 cDNA library fused with miRFP | used to construct stably express Sec61b-miRFP cell line in Caco-2 | This study |
| plko.1 | a lentiviral vector commonly used for RNA interference (RNAi) through stable expression of short hairpin RNAs (shRNAs). | Backbone plasmid | lab store |
| plko.1-shPFN1 | plko.1 derived plasmid containing PFN1 shRNAs predicted in website | used to construct stably knock down PFN1 cell line in Caco-2 | This study |
| plko.1-shFLNA | plko.1 derived plasmid containing FLNA shRNAs predicted in website | used to construct stably knock down FLNA cell line in Caco-2 | This study |
| plko.1-shCAP1 | plko.1 derived plasmid containing CAP1 shRNAs predicted in website | used to construct stably knock down CAP1 cell line in Caco-2 | This study |
| plko.1-shVASP | plko.1 derived plasmid containing VASP shRNAs predicted in website | used to construct stably knock down VASP cell line in Caco-2 | This study |
| plko.1-shDIAPH1 | plko.1 derived plasmid containing DIAPH1 shRNAs predicted in website | used to construct stably knock down DIAPH1 cell line in Caco-2 | This study |
| pLentiCRISPRv2 | a single-vector lentiviral CRISPR-Cas9 system developed for efficient gene knockout in mammalian cells. | Backbone plasmid | lab store |
| pcDNA3.1-sgRNA scaffold-U6 | a plasmid developed for loading dual sgRNAs | used to load dual sgRNAs targeting ITGA5 on sgRNA scaffold-U6 fragment | This study |
| pLentiCRISPRv2-dual ITGA5-sgRNAs | pLentiCRISPRv2 derived plasmids containing dual ITGA5 sgRNAs in structure as "U6-sgRNA1-sgRNA scaffold-U6- | used to knock out ITGA5 in Caco-2 cells | This study |

| | | | |
|---|---|---|---|
| | sgRNA2-sgRNA scaffold" | | |
| pcDNA3.1 | a commercially available mammalian expression vector widely used for high-level, constitutive expression of recombinant proteins in eukaryotic cells. | Backbone plasmid | lab store |
| pcDNA3.1-strep-ACTB | pcDNA3.1 derived plasmid containing strep-tag fused with ACTB cloned from Caco-2 cDNA library | used to express strep-tagged ACTB in 293F | This study |
| pET28a | a bacterial expression vector used for high-level production of recombinant proteins in *Escherichia coli*. | Backbone plasmid | lab store |
| pET28a-6His-ACD-Flag | pET28a derived plasmid containing ACD domain fused with 6His-tag in N-terms and Flag-tag in C-terms | used to express 6His-ACD-Flag in BL21 | This study |

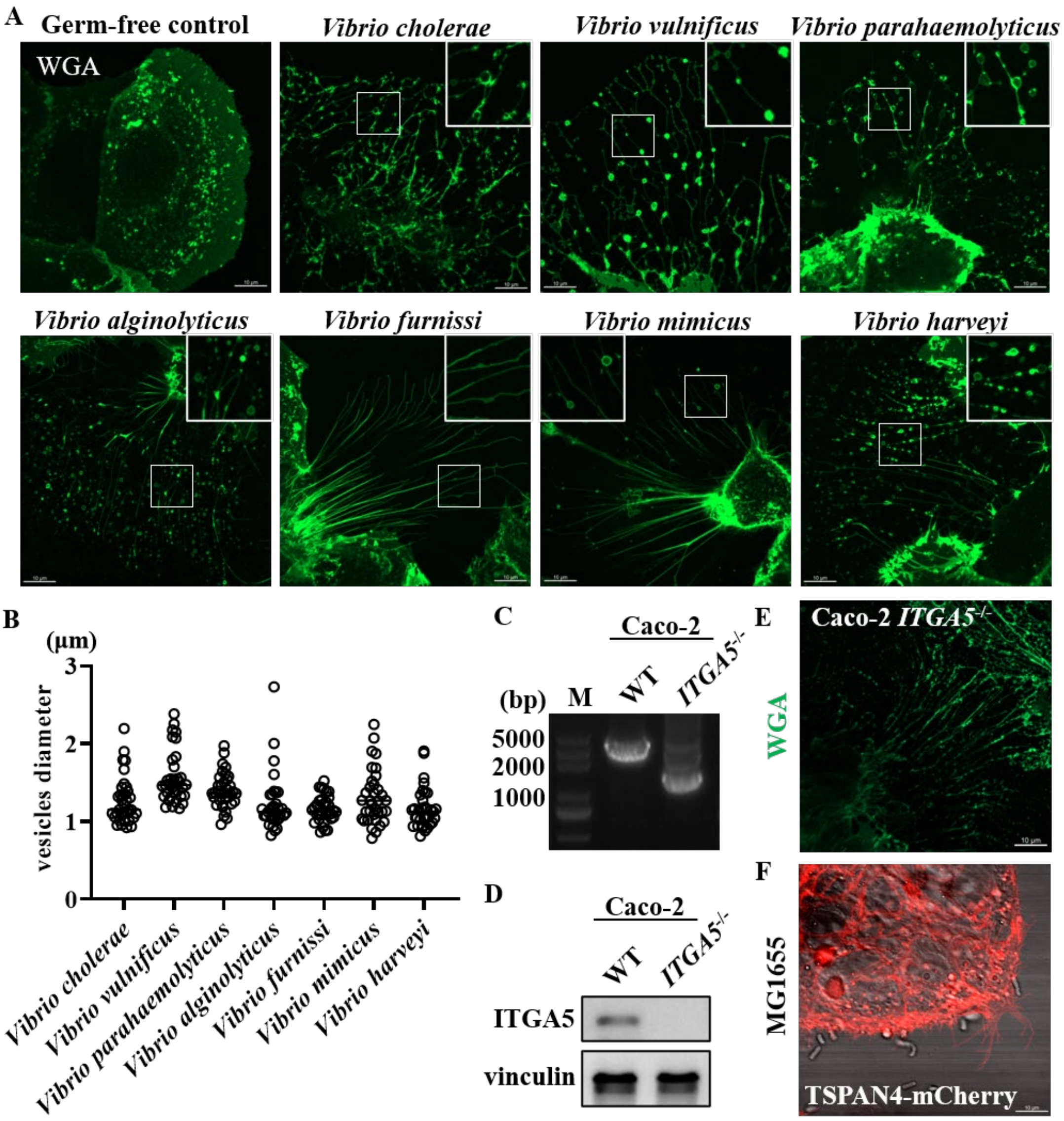


**Figure S1. *Vibrio* infection triggers morphological remodeling and vesicle formation in Caco2 cells.** (A)Wheat Germ Agglutinin (WGA) stained Caco-2 cells were infected by *Vibrios*. WGA is used to visualize plasma membrane to highlight morphological deformation, retraction fibers, and associated extracellular vesicles. (B) dimeters of the vesicles from (A) are measured as mean ± SEM; n=35. *P* values were calculated using one-way ANOVA followed by Dunnett's test. (C) Dual-sgRNA CRISPR/Cas9 approach was used to knockout *ITGA5*, and successful deletion was verified via PCR. (D) western blot assay confirming the loss of ITGA5 protein expression in *ITGA5*$^{-/-}$ cell lines. (E) Confocal images WGA stained Caco-2 defective express *ITGA5*. (F) Confocal images *E. coli* MG1655 infects Caco-2 cells express TSPAN4-mCherry.

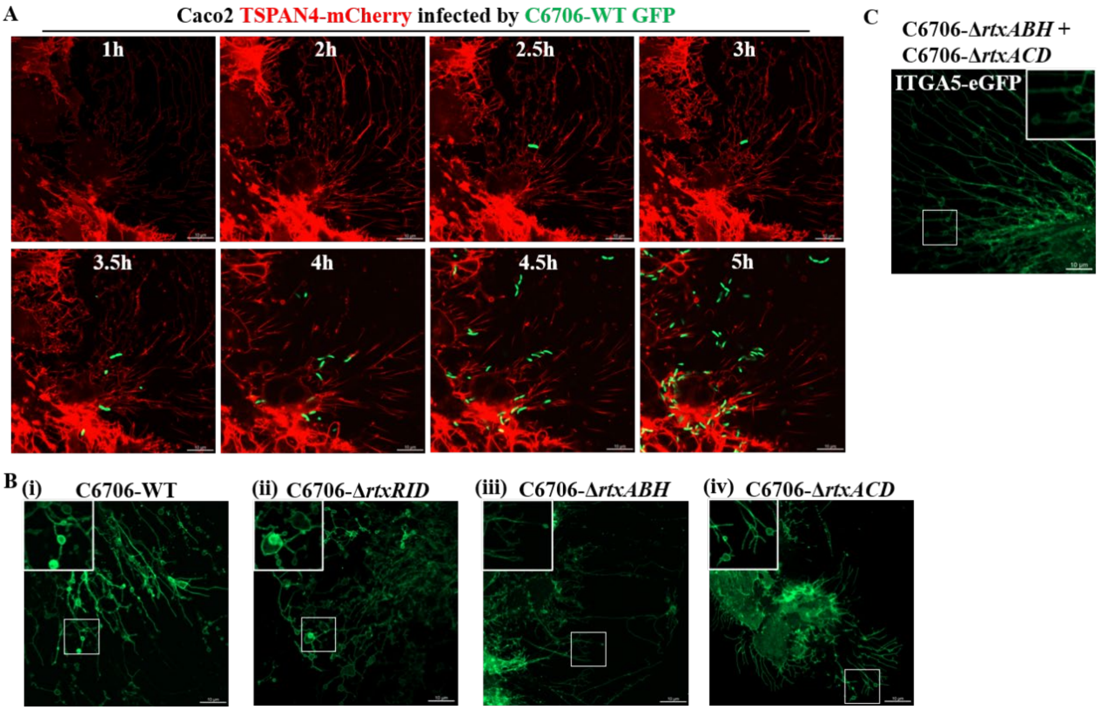


**Figure S2.** ***Vibrio*** **secretion products are sufficient to induce host cell passive migration.** (A) Caco2 stably express TSPAN4-mCherry and C6706 stably express eGFP. (B) representative confocal images Caco-2 stably express ITGA5-eGFP is infected by the indicated MARTX domain mutants. (C) confocal images Caco-2 infect with the mixture of C6706-Δ*rtxABH* and C6706-Δ*rtxACD*.

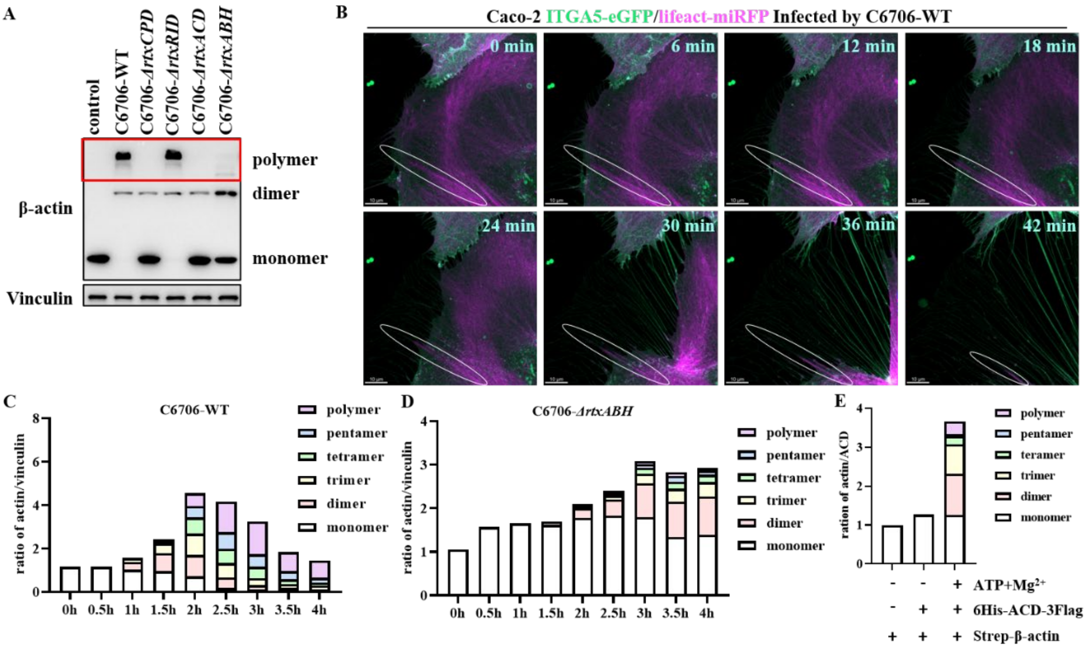


**Figure S3. ACD and ABH domains are essential for infection-induced actin crosslinking.** (A) Western blot analysis of high-order actin polymerization in Caco-2 cells. Samples were collected at 4 hours post-infection (MOI = 10). Vinculin was utilized as a loading control. (B) Time-lapse confocal images of Caco-2 cells stably expressing ITGA5-eGFP and Lifeact-miRFP following infection with C6706-WT. (C)-(E) Densitometric analysis of actin species distribution using ImageJ. The relative abundance of different actin oligomerization states was quantified from the Western blots presented in Figures 3A, 3C, and 3E.

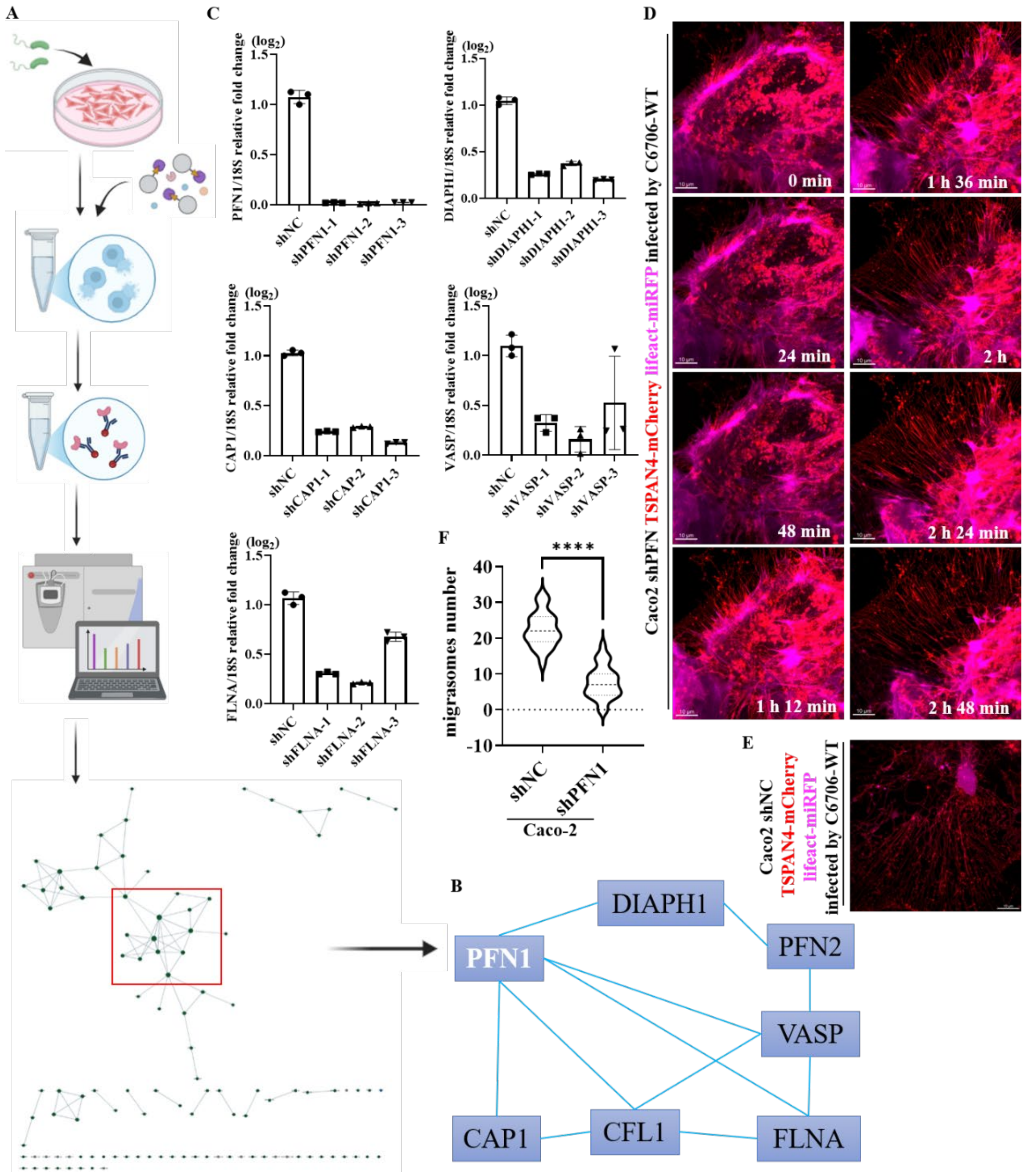


**Figure S4. IP-MS identifies actin modulators may function in migracytosis** (A) Schematic model of co-immunoprecipitation followed mass spectrum in Caco-2 infected with C6706-WT. β-actin was used as bite. (B) manually draw the cluster in (A). (C) qPCR reflect the knock down efficiency of the cluster proteins in (B) by shRNA. (D) Time-lapse confocal images of PFN1 knocked down Caco-2 cells stably expressing TSPAN4-mCherry and lifeact-miRFP following infection with C6706-WT. (E) Confocal images of negative control knocked down Caco-2 cells stably expressing TSPAN4-mCherry and lifeact-miRFP following infection with C6706-WT. (F) Quantification of the numbers of migrasome from (D) and (E) are measured as mean ± SEM; n=23-27. *P* values were calculated using t-test.

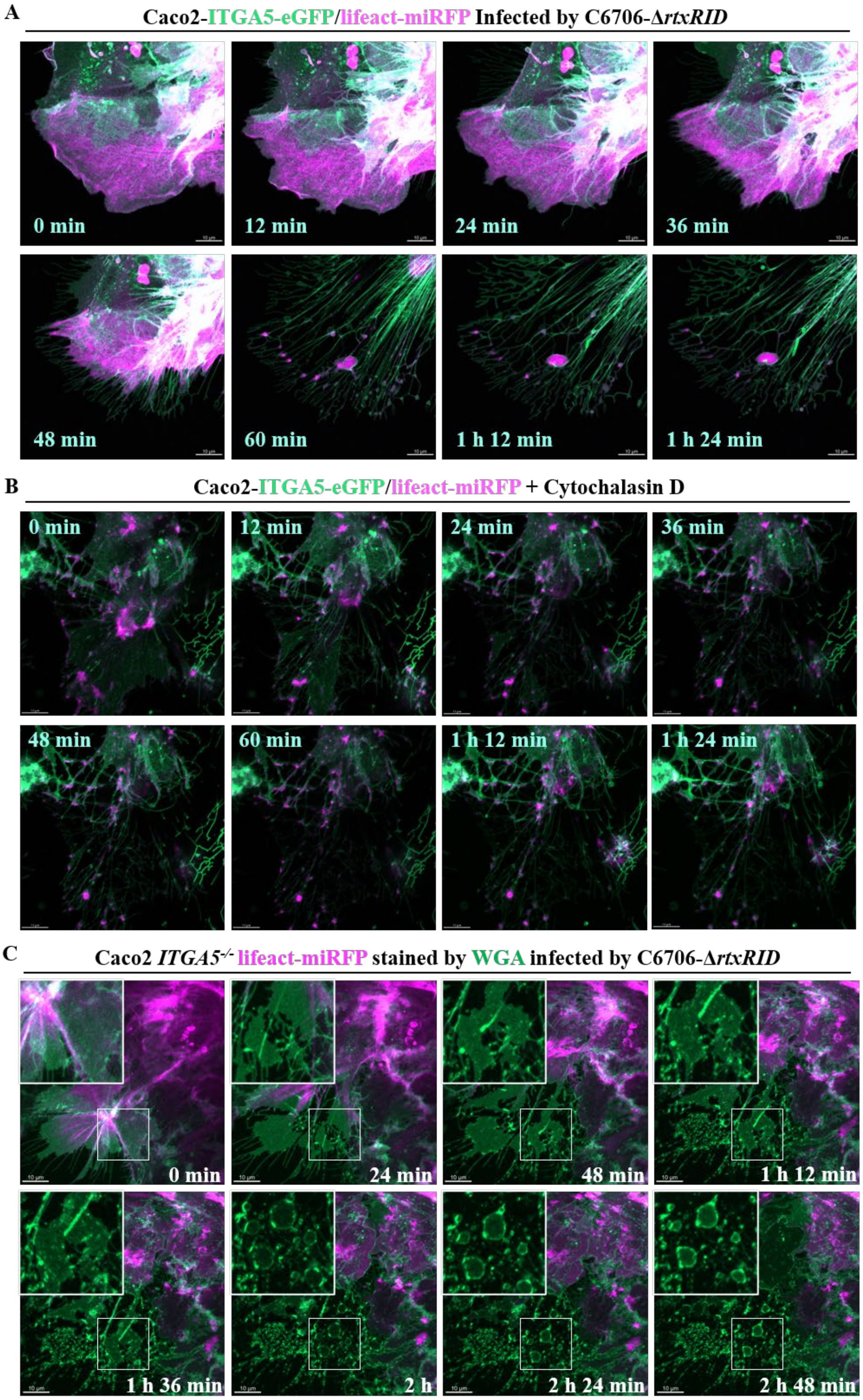


**Figure S5. Comparison of peripheral membrane sequestration in C6706-Δ*rtxrid* infection and Cytochalasin D treatment.** (A) and (B) Representative time-lapse confocal images Caco-2 cells stably expressing ITGA5-eGFP and Lifeact-miRFP. Cells were either (A) infected with the C6706-Δ*rtxRID* or (B) treated with 5 μM Cytochalasin D. (C) Time-lapse confocal images WGA stained Caco-2 *ITGA5*$^{-/-}$ cells stably expressing Lifeact-miRFP infected with the C6706-Δ*rtxRID.*

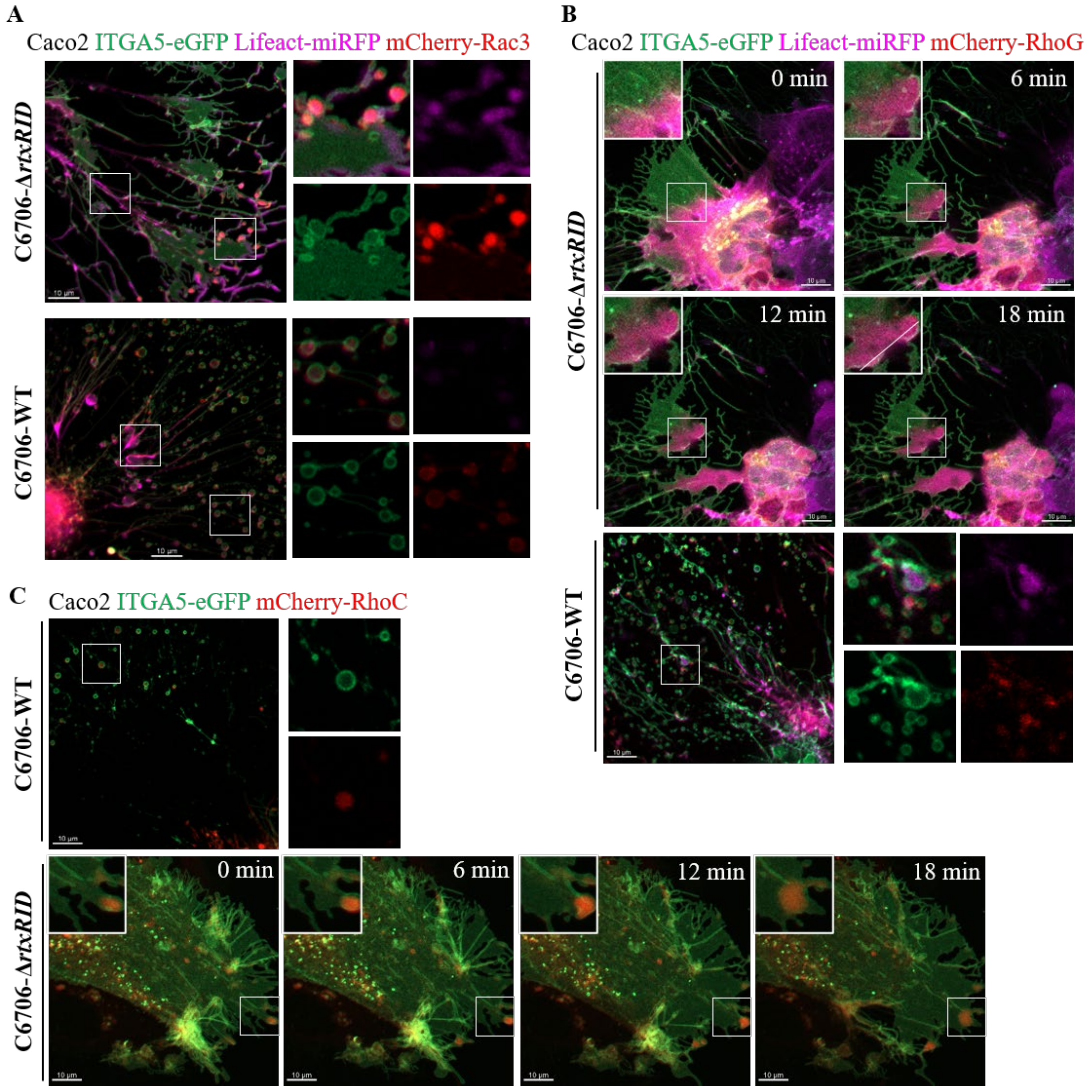


**Figure S6. Potential roles of Rho/RAS GTPases in stabilizing actin organization to rupture membrane during cell retraction.** (A)-(C) Live Caco-2 cells infected with either C6706-WT or C6706-Δ*rtxRID* were imaged by confocal time-lapse microscopy. (A)-(B) Cell stably expressing ITGA5-eGFP, Lifeact-miRFP and either mCherry-RAC3 (A) or mCherry-RhoG (B). (C) Cell stably express ITGA5-eGFP and mCherry-RhoC

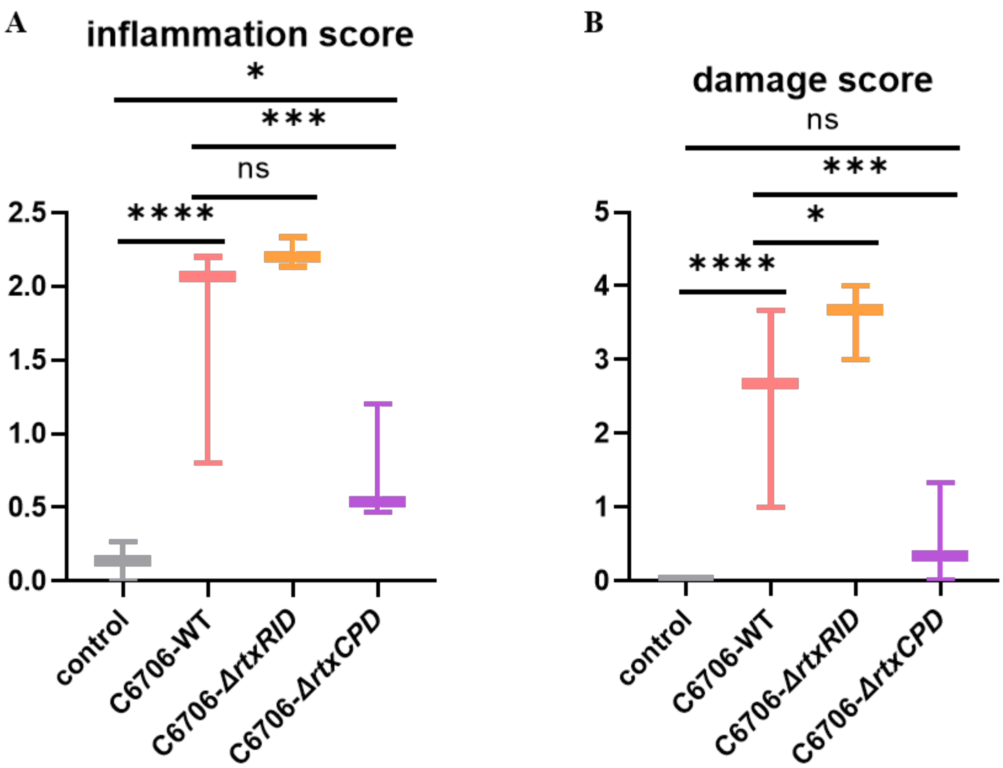


**Figure S7 Quantitative histopathological scoring of ileal sections from mice infected with *V. cholerae* strains.** (A) Composite inflammation score. (B) Epithelial damage score. Each point represents one animal (n=4 per group). P values were calculated using one-way ANOVA followed by Dunnett's test.
